# Oxygen-independent, hormonal control of HIF-1α regulates the developmental and regenerative growth of cardiomyocytes

**DOI:** 10.1101/2022.01.31.478572

**Authors:** Amarylis C.B.A. Wanschel, Angeliki Daiou, Jeffim N. Kuznetsoff, Stefan Kurtenbach, Katerina Petalidou, Thomai Mouskeftara, Helen G. Gika, Daniel A. Rodriguez, Eleftherios I. Papadopoulos, Georgios Siokatas, Polyxeni P. Sarri, Kyriaki Karava, Efthimios Tsivoglou, Christine Kottaridi, Alessandro G. Salerno, Krystalenia Valasaki, Wayne A. Balkan, Derek M. Dykxhoorn, Andrew V. Schally, Dimitrios L. Kontoyiannis, Antigone Lazou, Konstantinos E. Hatzistergos, Joshua M. Hare

## Abstract

Here, we present an O_2_-independent/HIF-1α (Hypoxia inducible factor-1α)-dependent mechanism that regulates developmental and regenerative growth of mammalian cardiomyoblasts. An autocrine feedback mechanism of GH/IGF1/SST (Growth hormone/Insulin-like growth factor 1/Somatostatin) signaling, mediated by GHRH/GHRH-R (Growth hormone-releasing hormone/GHRH-Receptor), is established specifically in NKX2-5 (NK2 Homeobox 5) expressing myocardial cells which affects HIF-1α stability through cAMP (cyclic adenosine monophosphate) or cGMP (cyclic guanosine monophosphate) activity. cAMP-mediated HIF-1α stabilization fuels Warburg metabolism and enhances NKX2-5 expression, limiting the developmental and regenerative growth of cardiomyoblasts. In contrast, cGMP-mediated HIF- 1α inhibition (or knock-out of HIF-1α) redirects glycolytically derived citrate toward long-chain saturated fatty acid biosynthesis, leading to enhanced developmental and regenerative growth of cardiomyoblasts. These findings suggest that HIF1α-mediated glycolysis serves as a rate-limiting, O_2_-independent sensor of cardiomyogenesis and that targeting GHRH/GHRH-R signaling could be a therapeutic strategy for regenerating the mammalian heart post-injury.

**Summary Figure.**
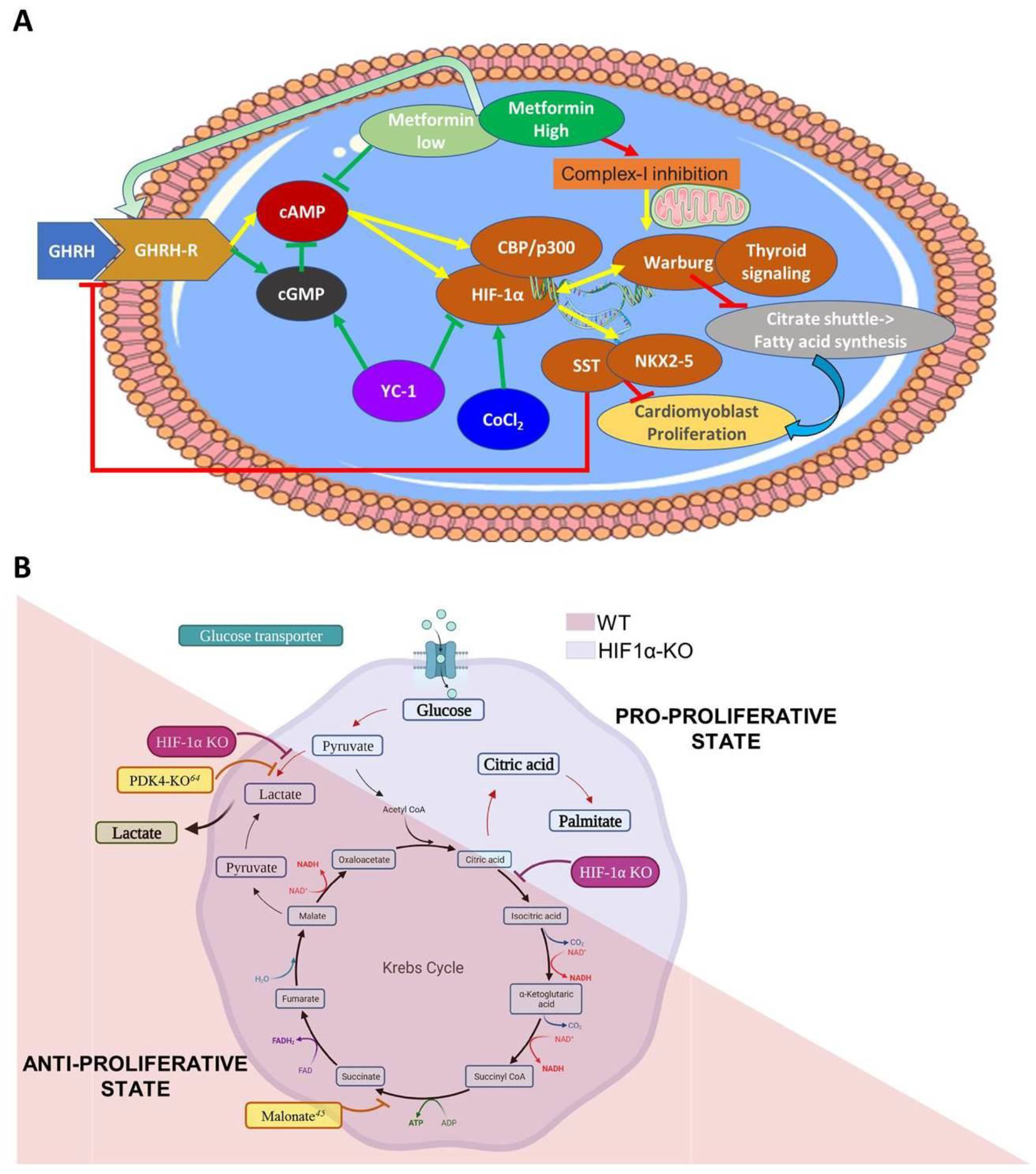
Cardiomyoblast proliferative capacity is regulated by HIF1α-mediated metabolic profile, through an NKX2-5 lineage-specific, O_2_-independent feedback mechanism. **(A)**, HIF-1α is aerobically regulated through the intracellular activation of cAMP and cGMP in response to a GH1/IGF1/SST feedback mechanism, which activates the GHRH/GHRH-R signaling pathway in NKX2-5^+^ cardiomyoblasts. Stabilized HIF-1α (e.g. by CoCl_2_) promotes Warburg metabolism and cell cycle exit, whereas knockout or inhibition (e.g. YC-1) of HIF-1α or cAMP (low-dose metformin) promotes *de novo* fatty acid synthesis and enhances cardiomyoblast proliferation. **(B)**, HIF-1α aerobic stabilization switches the metabolism towards glycolysis and restricts cardiomyoblast proliferation. Contrariwise, knockout of HIF-1α promotes oxidative metabolism in the Krebs cycle rather than lactate synthesis, thus stimulating the proliferative and regenerative capacity of cardiomyoblasts.

## Introduction

The human heart develops early during embryogenesis to produce the four chambered mammalian heart capable of efficiently moving blood throughout the body. The evolutionary enhancements that have made the four chambered heart such an effective tissue are offset by the lack of regenerative potential seen in amniotes, of which humans are representative, compared to anamniotes^1^. These evolutionary adaptations are linked to the hormonal and metabolic mechanisms that contributed to the rise of sexual dimorphism and endothermy^2^. For example, hermaphroditic sea slugs can repeatedly autotomize at their neck region and simultaneously regrow an entire new body, including a whole new functional heart, out of the remaining head tissue^3^. Zebrafish^4^ and some urodele amphibians (e.g. salamanders)^5^ retain life-long cardiac regenerative capacity in response to injury; whereas, anurans (frogs and toads) lose cardiac regenerative potential after metamorphosis^1^. In mammals, the cardiac regenerative window closes a few days after birth^6^. Due to their relatively long life, this lack of regenerative capacity has rendered humans particularly vulnerable to heart disease^7^. Therefore, understanding human developmental and regenerative mechanisms of cardiac myogenesis could provide important insights for developing novel, cardiac regenerative therapies.

Studies in model organisms with lifelong or transient cardiac regenerative capacity indicate that heart regeneration is orchestrated through an O_2_-dependent metabolic rewiring of the damaged tissue which stimulates epimorphic growth^1^. In the “solar-powered” sea slug, this is achieved through kleptoplasty, whereby the regenerating cells of the autotomized head phagocytose chloroplasts from the surrounding algae to bioengineer an O_2_-rich intracellular environment that fuels, through photosynthesis, the energy demands required to support organismal survival and *de novo* heart regeneration^3^. Interestingly, the O_2_-dependent metabolic rewiring in the “carbon-powered” vertebrate heart seems to operate in the opposite manner. Specifically, O_2_-rich environments are detrimental to both development and regeneration in zebrafish^8^ and neonatal mouse hearts^9^. Consequently, enhancing anaerobic over aerobic metabolism through either the stabilization of the O_2_-sensing transcription factor Hypoxia-inducible factor (HIF)-1α^8,10^ or promoting NRG1/ErbB2 signaling^11^, stimulates cardiomyocyte proliferation and regeneration in zebrafish and mice.

However, the roles of O_2_ and its molecular sensor HIF-1α in heart development and regeneration are not well understood. For example, Iyer and colleagues demonstrated that the loss of HIF-1α in mice results in reduced glucose fermentation promoting excessive myocardial proliferation in the embryonic heart, such that the fetal chambers are filled up with cardiomyocytes by E10.5 resulting in death^12^. Expanding on this study, several HIF-1α^–/–^ mutant mice have been generated that confirmed the role of HIF-1α as a master transcriptional regulator of cardiac glycolytic metabolism; however, the cardiac phenotypes have been surprisingly inconsistent; i.e., impaired glycolytic metabolism in response to HIF-1α knockout, has resulted in normal^13,14^, hypoplastic^15,16^, and hyperplastic^12,17^ myocardial phenotypes. Thus, although most findings agree on a fundamental, evolutionary role of an O_2_-dependent metabolic rewiring for the growth of new myocardial tissue in the developing and regenerating heart, is unknown whether a HIF1α-mediated glycolytic rewiring acts as a “make”^9,11,16,18^ or “break”^3,12,19^ signal in mammals.

Therefore, we sought to directly address the role of HIF-1α in mammalian cardiomyocyte proliferation and regeneration. To that end, we knocked-out HIF-1α (HIF1α-KO) in human induced pluripotent stem cell (hiPSCs) lines and analyzed their cardiomyocyte differentiation and proliferation capacity through stage-specific, multiomic profiling. In addition, we manipulated pharmacologically HIF-1α stability and glucose metabolism in neonatal mice undergoing heart regeneration. Our findings provide important new insights that establish the role of HIF-1α-mediated glycolysis as a rate-limiting sensor of human cardiomyocyte proliferation. Importantly, we demonstrated that myocardial HIF-1α signaling is regulated through an oxygen-independent mechanism involving autocrine stimulation of the growth hormone-releasing hormone (GHRH) signaling pathway specifically in NKX2-5-expressing cardiomyocytes. Hence, targeting the GHRH signaling pathway could serve as a novel therapeutic strategy for specifically modulating HIF-1α activity in human cardiomyocytes to stimulate their proliferative growth following injury.

## Results

### HIF-1α limits cardiomyogenesis in humans

The various HIF-1α^–/–^ mouse models have produced inconsistent phenotypes that have obscured the role of HIF-1α signaling in cardiac development and regeneration^1,12-16,20,21^. To overcome these inconsistencies and examine the role of HIF-1 □ in human cells, we generated two HIF1α-KO hiPSCs lines using CRISPR/spCas9-based genome editing^22,23^ [Suppl. Fig. 1A]. Compared to the current experimental models, the HIF1α-KO hiPSCs approach offers several advantages. Firstly, hiPSCs can be subjected to highly efficient, chemically defined, guided differentiation into cardiomyocytes, thereby providing tissue-specific modeling capabilities with minimal confounding effects secondary to extracardiac HIF-1α mechanisms. Secondly, the global HIF-1α knock out used in this model can minimize any tissue- or stage-specific inconsistencies, such as those observed in response to Cre/LoxP-mediated HIF-1α^–/–^ mice^1^. Thirdly, the system is human and, therefore, eliminates any potential species-specific evolutionary differences that may govern cardiac HIF-1α signaling.

Both HIF1α-KO hiPSC lines exhibited loss of HIF-1α protein expression and could be propagated in E8 medium at 37°C and 5% CO_2_, without obvious phenotypic differences compared to wild-type (WT) parent cells [Suppl. Fig. 1B-D]. Importantly, both HIF1α-KO lines exhibited similar cardiomyocyte differentiation capacities, and therefore have been used interchangeably throughout the study [Suppl. Fig. 1E]. Intriguingly, during the phenotypic characterization, we observed that HIF1α-KO hiPSCs generated beating cardiomyocytes more consistently, compared to their WT hiPSCs counterparts [Suppl. Video 1]. To gain further insights, WT and HIF1α-KO hiPSC cardiomyocytes were subjected to next-generation RNA sequencing profiling (RNA-seq). A total of 27,868 protein-coding and non-coding genes were found to be expressed in both WT and HIF1α-KO cardiomyocytes [Suppl. Data 1]. Of these, 1,445 genes were significantly down-regulated, and 1,267 genes were significantly up-regulated in HIF1α-KO compared to WT cardiomyocytes (q-val<0.05), [Suppl. Data 1 and Fig. 1A], which is consistent with HIF-1α operating both as a transcriptional activator, as well as repressor in the developing human myocardium^16^. Importantly, since all experiments were performed under normal atmospheric oxygen levels, these findings indicate that HIF-1α operates in an O_2_-independent manner during human cardiomyogenesis. This result is consistent with previous *in vivo* experiments in mice, which show that HIF-1α activity becomes spatiotemporally compartmentalized in the developing mouse heart^14,16^; and this compartmentalization is unlikely to be O_2_-dependent because the staining patterns of the hypoxia reporter pimonidazole and nuclear HIF-1α, do not consistently overlap^24-26^ [Suppl. Fig. 2].

**Figure 1.**
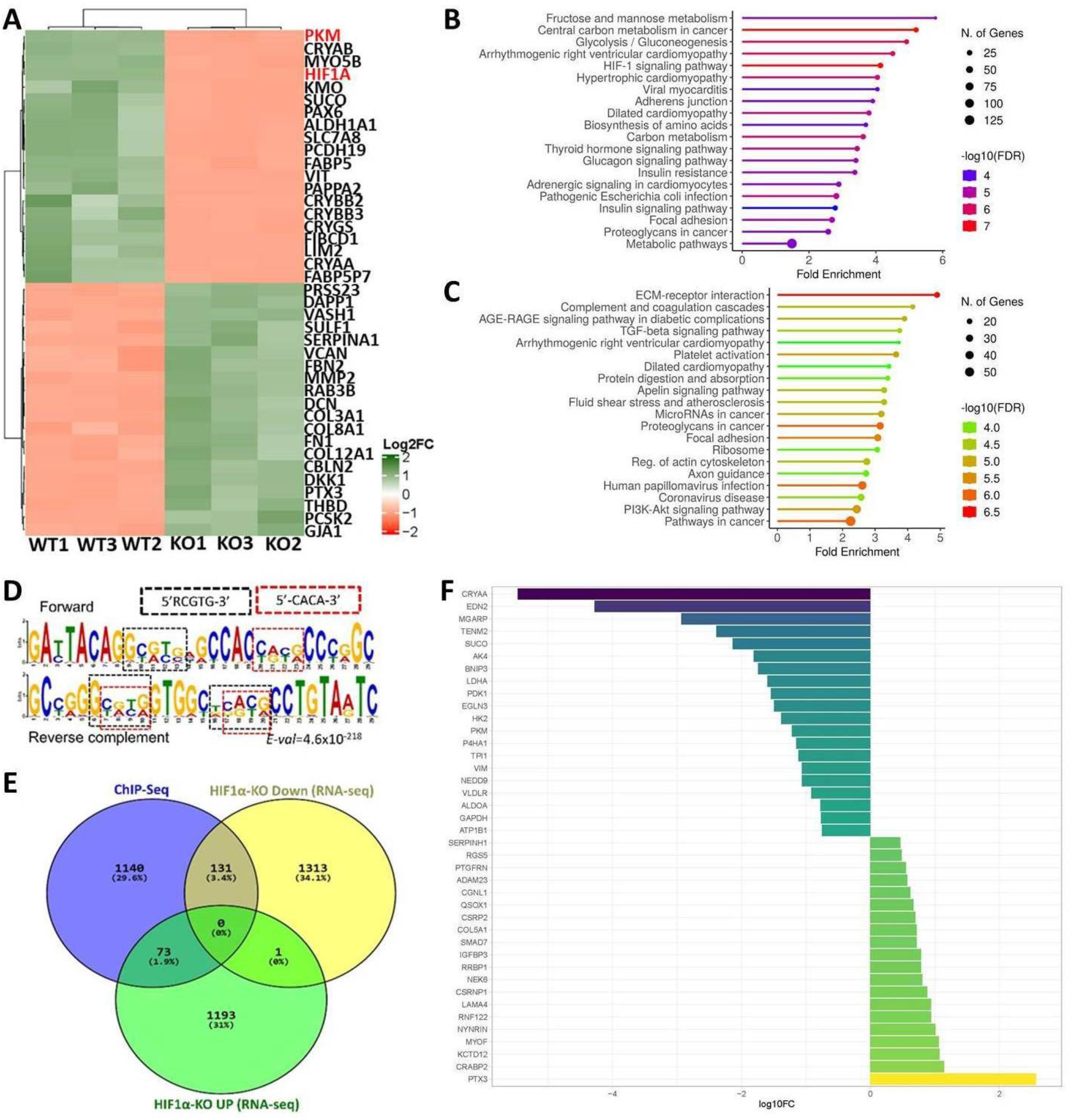
Determination of cardiac gene regulatory programs under the control of *HIF-1α* through combined CRISPR/Cas, RNA-seq and ChIP-seq analyses. **(A)**, Unsupervised hierarchical clustering of the top-20 up-and down-regulated genes (n=3 replicates/group). **(B)**, Lollipop plot of the top-20 KEGG terms enriched in 1,445 significantly downregulated genes in HIF1α-ΚΟ vs WT cardiomyocytes. **(C)**, Top-20 KEGG terms enriched in 1,267 significantly upregulated genes in HIF1α-ΚΟ vs WT cardiomyocytes. **(D)**, MEME motif of the DNA sequences ±50bp from the HIF-1α ChIP-seq peak summits with a q-val ≤ 1×10^−100^. **(E)**, Venn plot illustrating the overlaps between the RNA-seq (green, yellow) and ChIP-seq (blue) datasets (note a common gene in HIF-1a-KO UP and DOWN datasets corresponding to different transcript variants). **(F)**, Top-20 up- and down-regulated HIF-1α target genes.

KEGG functional enrichment analysis of the down-regulated genes showed significant overrepresentation in HIF-1α signaling, central carbon metabolism, as well as thyroid hormone signaling, arrhythmogenic right ventricular cardiomyopathy, hypertrophic cardiomyopathy, and dilated cardiomyopathy pathways [Fig. 1B and Suppl. Data 2]. In contrast, upregulated genes were functionally enriched in cell-ECM interaction/focal adhesion processes, as well as in AGE-RAGE and PI3K/AKT intracellular signaling pathways [Fig. 1C].

Next, ChIP-seq analysis was performed to determine which of the differentially expressed genes were directly regulated by HIF-1α. Accordingly, WT hiPSC-cardiomyocytes were treated with the hydroxylase inhibitor CoCl_2_ (100μM for 24h) to stabilize HIF-1α and ensure maximal DNA binding capacity. The HIF-1α bound chromatin was crosslinked, sonicated, immunoprecipitated using the previously KO-validated HIF-1α antibodies, and sequenced. We identified 1,416 peaks, corresponding to 1,344 genes, ∼57% of which were in promoter regions. Motif discovery analysis identified a motif containing 5’-RCGTG-3’ and 5’-CACA-3’ sequences (E-value= 4.6×10^−218^), closely resembling the hypoxia response element (HRE) motif to which HIF-1α binds [Fig. 1D]^27^. Alignment of the RNA-seq and ChIP-seq datasets revealed that only 131/1,445 and 73/1,267 significantly down- and up-regulatedd genes in the HIF-1□ KO cells campared to the HIF-1□ isogenic parental cells, respectively, were HIF-1αtargets [Fig. 1E and Suppl. Data 3], suggesting that >90% of the RNAseq-detected differentially expressed genes were indirectly regulated by HIF-1α.

KEGG functional enrichment analysis of the 131 genes that were both down regulated in the HIF-1α KO cells and shown to be targets of HIF-1α by ChIP seq showed significant overrepresentation in glucose metabolism and HIF-1α signaling, indicating that loss of HIF-1α in human myocardial cells significantly alters aerobic glycolysis, also known as the Warburg effect^28,29^ [Fig. 2A]. For example, *LDHA, PDK1, EGLN3, HK2*, and *PKM* were among the top glycolytic genes found to be directly bound and transcriptionally activated by aerobically stabilized HIF-1α in the human myocardium [Fig. 1F, Suppl. Data 4]. In parallel, genes that were indirectly activated in response to aerobically stabilized HIF-1α were functionally enriched in cardiomyopathy related processes, as well as in adrenergic and thyroid hormone signaling pathways [Fig. 2B and Suppl. Data 4]. Notably, increased thyroid hormone signaling has been evolutionarily linked to metabolic mechanisms related to the postnatal acquisition of endothermy and the loss of cardiac regenerative capacity in mammals^2^. In the HIF1α-KO hiPSC cardiomyocytes, thyroid hormone signaling genes that were indirectly repressed include the Nuclear Receptor Coactivators 1 and 2 (*NCOA1* and *NCOA2*), which play important roles in the regulation of the hypothalamic-pituitary-thyroid axis^30^; and the Type II iodothyronine deiodinase (*DIO2*) that controls T4 deiodination to the bioactive T3 form^31^.

**Figure 2.**
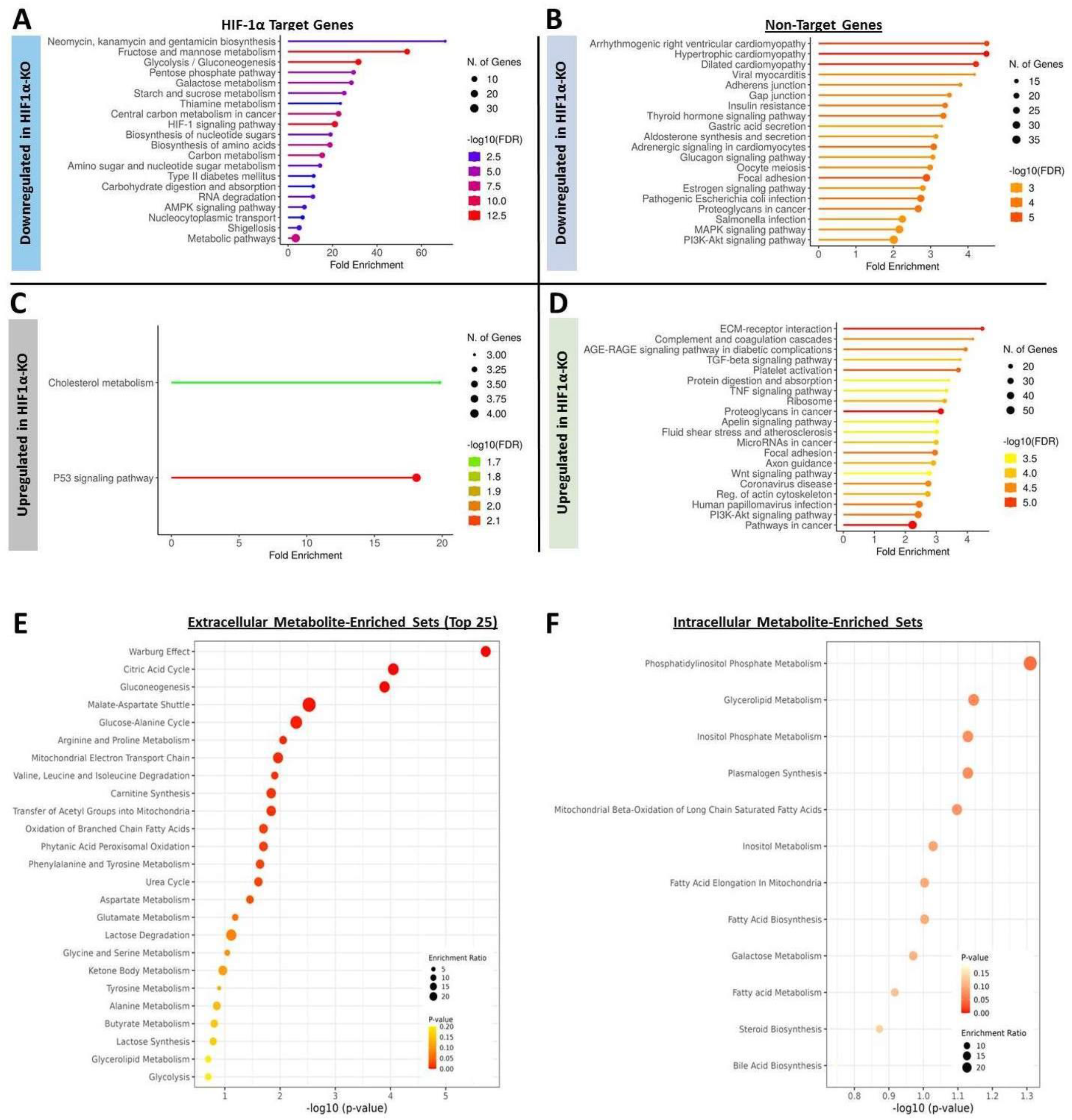
Functional enrichment analyses of HIF-1α target and non-target differentially expressed genes between wild type (WT) and HIF1α-ΚΟ (KO) human induced pluripotent stem cell (hiPSC)- derived cardiomyocytes. **(A)**, Top-20 KEGG terms enriched in the 131 HIF-1α target genes that are significantly downregulated in HIF1α-ΚΟ vs WT cardiomyocytes. **(B)**, Top-20 KEGG terms enriched in the 1,313 HIF-1α non-target genes that are significantly downregulated in HIF1α-ΚΟ vs WT cardiomyocytes. **(C)**, Two KEGG terms found to be significantly enriched in the 73 HIF-1α target genes that are significantly upregulated in HIF1α-ΚΟ vs WT cardiomyocytes. **(D)**, Top-20 KEGG terms enriched in the 1,193 HIF-1α non-target genes that are significantly upregulated in HIF1α-ΚΟ vs WT cardiomyocytes. **(E)**, Dot plot of top-25 Small Molecule Pathway Database (SMPDB)-enriched terms following over representation analysis of significantly different extracellular metabolites between WT and HIF1α-ΚΟ hiPSC-derived cardiomyocytes. **(F)**, SMPDB-enriched terms following over representation analysis of significantly different intracellular metabolites between WT and HIF1α-ΚΟ hiPSC-derived cardiomyocytes.

KEGG functional enrichment analysis of the 73 HIF-1α target genes that were upregulated in HIF1α-KO hiPSC cardiomyocytes illustrated significant overrepresentation in p53 signaling (*GADD45B, IGFBP3, PERP, SERPINE1*) and cholesterol metabolism *(LRP1, APOH, ANGPTL4*) pathways, suggesting that HIF-1α directly binds and suppresses these genes in human myocardial cells [Fig. 2C, Suppl. Data 4]. In parallel, genes that were indirectly repressed in response to aerobically stabilized HIF-1α were functionally enriched in ECM-receptor interaction related processes, as well as in AGE-RAGE and PI3K/AKT signaling pathways [Fig. 2D, Suppl. Data 4].

Given the strong direct and indirect (i.e. thyroid hormone signaling inhibition) effects of HIF-1α on the transcriptional regulation of cardiomyocyte metabolism, WT and HIF1α-KO hiPSC were subjected to gas chromatography-mass spectrometry (GC-MS) based analysis of intracellular and extracellular metabolites, both before and after cardiomyocyte differentiation^32^. A total of 37 intracellular and 29 extracellular metabolites were detected. Among these, statistical analysis detected 7/37 intracellular and 1/29 extracellular metabolites to differ significantly between undifferentiated (differentiation day 0) WT and HIF1α-KO hiPSCs [Suppl. Fig. 3A]. The strongest effect was observed in D-glucose levels which did not differ extracellularly but were significantly reduced intracellularly (∼2.3-fold decrease, *p*=0.0016) in WT relative to HIF1α-KO hiPSCs [Suppl. Fig. 3A]. These findings, in addition to increased intracellular levels in Krebs cycle intermediates (i.e. citrate and malate), indicate increased glycolysis in WT relative to HIF1α-KO hiPSCs through aerobic activation of HIF-1α. Notably, an important role for glycolysis in human pluripotent stem cells has been previously reported and thought to be largely MYC-dependent^33,34^.

In day 10 spontaneously beating WT and HIF1α-KO hiPSCs cardiomyocytes, there were a total of 3/37 intracellular and 11/29 extracellular metabolites that were significantly different [Suppl. Fig. 3B-C]. Metaboanalyst enrichment analysis^35^ of extracellular metabolites demonstrated significant overrepresentation in Warburg effect- and Krebs cycle-related processes [Fig. 2E]; whereas intracellular metabolites showed weak enrichment in inositol and glycerolipid metabolism [Fig. 2F].

Combined, the RNA-seq, HIF-1α ChIP-seq and metabolomic data demonstrate that, during cardiac myogenesis, HIF-1α is stabilized in an O_2_-independent manner to activate a glycolytic gene program that leads to fetal cardiomyocyte overflow metabolism (Warburg effect), as indicated by the elevated lactate/decreased glucose levels and accumulation of Krebs cycle intermediates in the extracellular space of WT, but not HIF1α-KO hiPSC cardiomyocytes. In addition, the increased intracellular saturated long-chain fatty acid levels in HIF1α-KO hiPSC cardiomyocytes suggest that induction of aerobic glycolysis through HIF-1α promotes oxidation and extracellular leakage of citrate, prohibiting its shuttling toward fatty acid biosynthesis^36^ [Suppl. Fig. 3B-C]. Moreover, because palmitate inhibits thyroid hormone signaling^37^, the increased intracellular palmitate concentration corroborates the indirect effects of HIF-1α on thyroid hormone signaling genes [Fig. 2B].

We next examined the consequences of HIF-1α knock-out on cell proliferation during human cardiomyogenesis. Time-resolved cell-cycle analysis using propidium iodide-based flow cytometry confirmed that the loss of HIF-1α is associated with reduced proliferative capacity of hiPSCs, as indicated by the higher fraction of cells in G2/M phase in WT relative to HIF1α-KO hiPSCs [Fig. 3A-B]. This finding agrees with the positive role of hypoxia in human pluripotent stem cell proliferation^38^. However, upon the onset of cardiomyocyte differentiation, the cell cycle profiles reversed, with HIF1α-KO hiPSCs exhibiting increasingly higher fractions of G2/M phase cells, relative to their WT counterparts. The differences in cell cycle activity peaked on day 7 of differentiation, with HIF1α-KO hiPSCs yielding 36.8%±13.5% more cardiomyoblasts in G2/M phase, relative to WT (*p*= 0.03) [Fig. 3A-B]. Confocal immunofluorescence analysis for Ser-10 phosphorylated histone H3 (PH3) confirmed a ∼2.15-fold higher mitotic activity (p=0.0003) in day 7 HIF1α-KO vs. WT hiPSC cardiomyoblasts [Fig. 3C-D]. More importantly, co-staining with HIF-1α demonstrated strong nuclear immunoreactivity in day 7 WT cardiomyoblasts which, however, never co-localized with PH3 nuclei [Fig. 3C-D]. In addition, immunoreactivity of nuclear HIF-1α continued to increase on day 10 WT cardiomyoblasts [Suppl. Fig. 1E], and this effect was accompanied by reduced mitosis [Fig. 3D]. These findings support the hypothesis that aerobic stabilization of HIF-1α is a rate-limiting mechanism for human cardiomyoblast proliferation.

**Figure 3.**
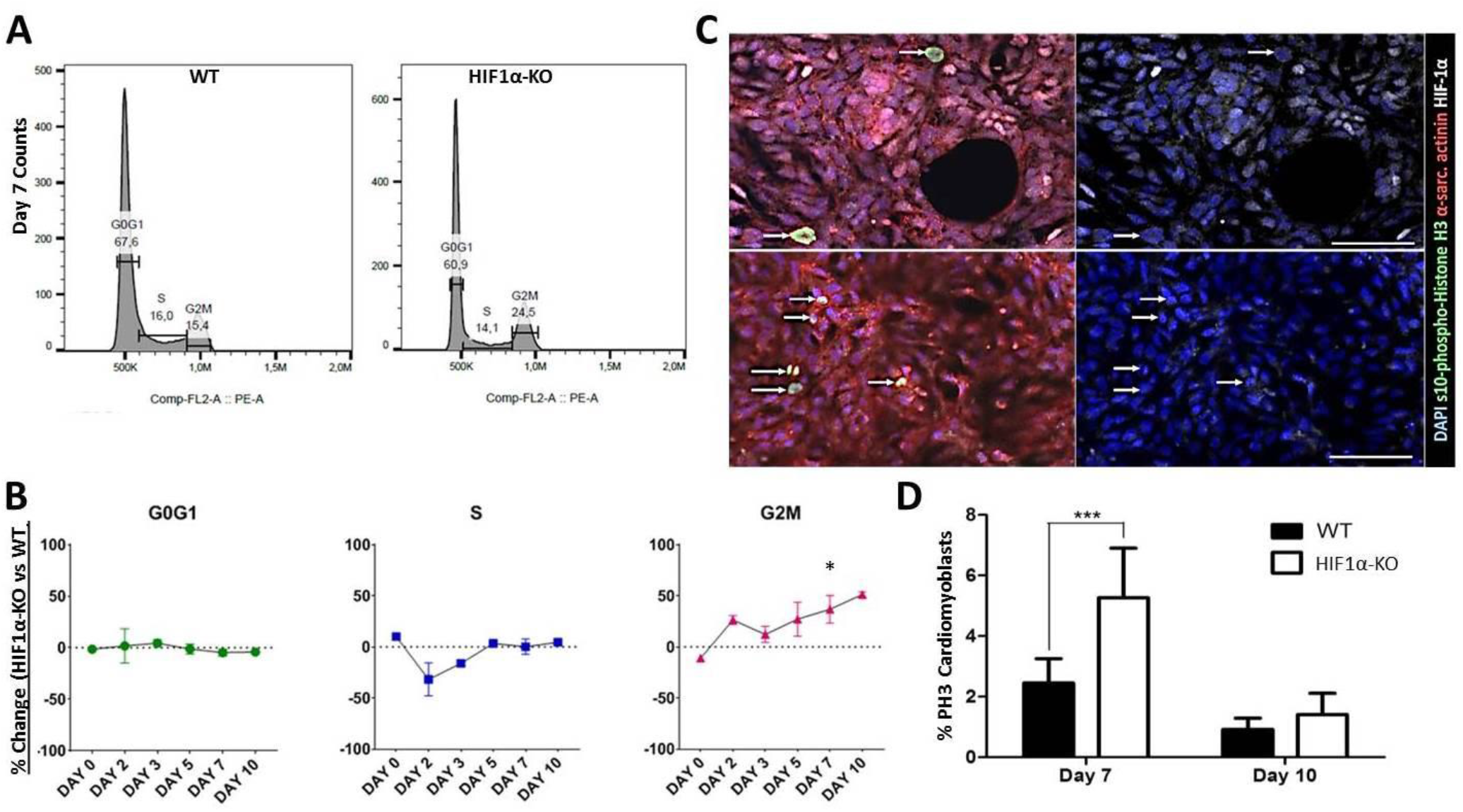
HIF-1α limits cell proliferation during human cardiomyogenesis. **(A)**, representative flow cytometry histograms following cell cycle analysis with propidium iodide, during hiPSC cardiomyocyte differentiation. **(B)**, Line graphs illustrating temporally resolved % changes in cell cycle phases (G0/G1, S and G2/M) in HIF1α-ΚΟ relative to WT hiPSC derivatives. **(C)**, confocal microscopy on day 7 of differentiation, demonstrating lack of nuclear HIF-1α immunoreactivity in both WT (upper panels, arrows) and HIF1α-KO mitotic cardiomyoblasts (lower panels, arrows). Mitotic cardiomyoblasts are defined as ser10-phospho-Histone H3^+^/α-sarcomeric actinin^+^ cells. **(D)**, Confocal immunofluorescence quantification of mitotic cardiomyoblasts. (*p<0.05, ***p<0.001). (n=2-3/days 0, 2, 3, 5, and 10; n=6/day 7).

### HIF-1α inhibition enhances cardiomyocyte regeneration in neonatal mice

In mammals, cardiomyogenesis ceases before birth, but for a short period during the first week of life cardiomyocytes retain the capacity for regeneration^6^. To test whether signaling of HIF-1α is also involved in cardiomyocyte regeneration, we employed a neonatal mouse model of cryoinjury-induced heart regeneration^39^. Accordingly, 1 day old BALB/c mice were randomized to sham or cryoinjury operation, followed by daily subcutaneous injections of PBS (control), the HIF-1α inhibitor/soluble guanylyl cyclase (sGC) agonist YC-1^40^ (70μg/50μL), or the HIF-1α inducer CoCl_2_ (120μg/50μL)^41^. Animals were followed for 7 days, a time-point previously associated with the peak in proliferative activity of regenerative cardiomyocytes^6^. The overall survival rate was 78.3% for cryoinjured vs 85.7% for sham, with no significant differences between groups [Suppl. Fig. 4A-B]. As expected, HIF-1α exhibited robust nuclear immunoreactivity in CoCl_2_-treated animals and virtually complete loss in HIF-YC-1 treated, animals [Suppl. Fig. 4C]. In addition, in all the studied groups, subcutaneous injection of pimonidazole three hours prior to euthanasia (Hypoxyprobe, 60mg/kg) did not produce positive immunostaining, indicating an absence of hypoxia in the day 7 post-cryoinjury myocardium [Suppl. Fig. 4C]. Moreover, compared to other groups, CoCl_2_ treated hearts exhibited increased sphericity [Suppl. Fig. 4B]. This observation phenocopies transgenic mice that have NKX2-5 cardiomyoblast-specific VHL ablation that has been attributed to abnormalities of HIF-1α-induced conduction system ^14^. Finally, confocal immunofluorescence analysis examining PH3 demonstrated that, compared to the control group, YC-1-treated hearts exhibited an ∼25% increase in mitotically active, regenerative cardiomyocytes; whereas CoCl_2_-mediated stabilization of HIF-1α attenuated cardiomyocyte proliferation by ∼10% [Fig. 4A-B].

**Figure 4.**
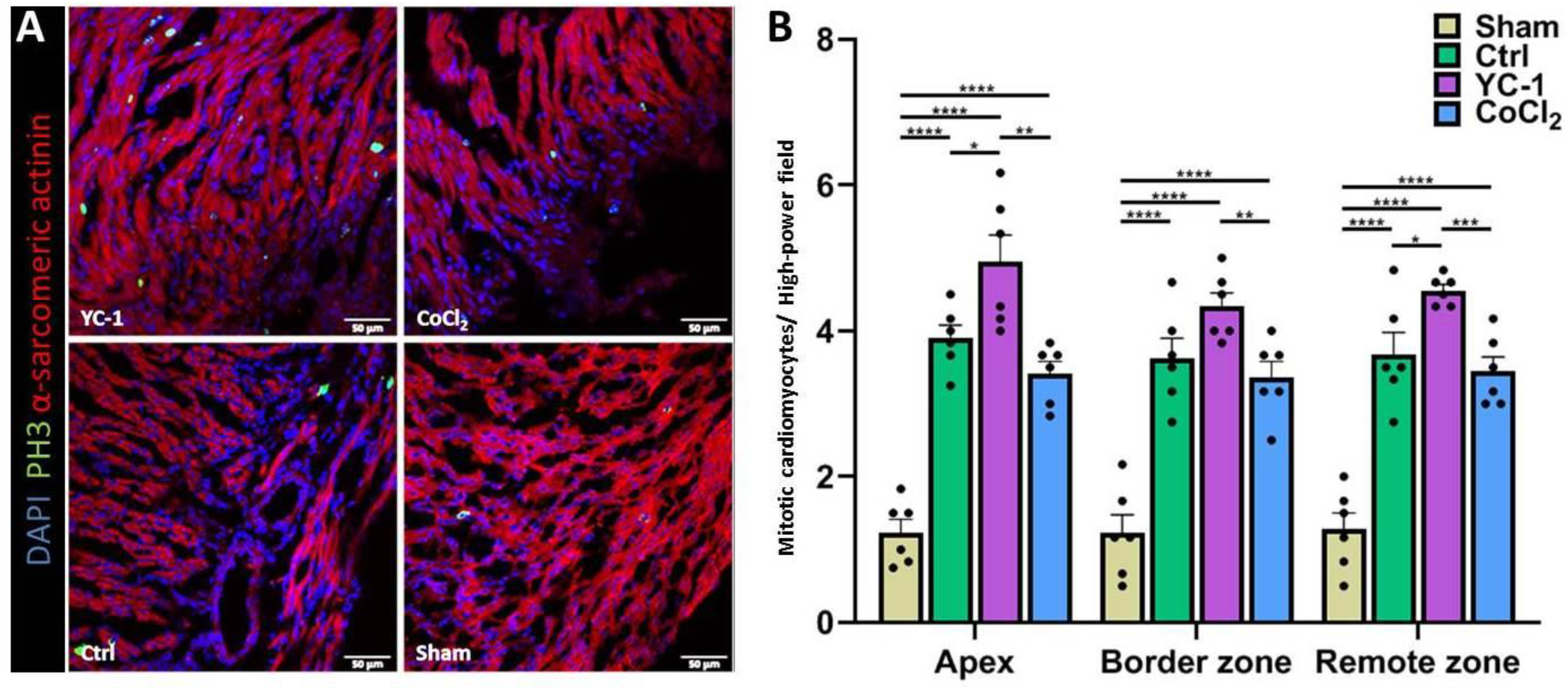
HIF-1α limits cardiomyocyte proliferation after cryoinjury. **(A)**, representative confocal microscopy images from the apical region of the heart of mice that received YC-1, CoCl2, or saline after surgery with or without cryoinjury. **(B)**, cardiomyocyte proliferation is induced after myocardial injury. Inhibition of HIF-1α by subcutaneous administration of YC-1 increases the number of mitotic cardiomyocytes, whereas its stabilization by administration of CoCl_2_ decreases the proliferation levels of cardiac myocytes. PH3, ser10-phospho-Histone H3. Ctrl, control. Scale bars, 50μm. (***p≤0.001, **p≤0.01, *p≤0.05), (n=6/group).

These findings are consistent with previous work suggesting activation of HIF-1α in the postnatal and regenerating vertebrate heart^8,10,42^, but extend these earlier findings to show that activation of HIF-1α occurs in an O_2_-independent manner to restrict, rather than promote, cardiomyocyte proliferation in the regenerating neonatal mouse heart.

Since regeneration of vertebrate heart has been paradoxically linked to HIF-1α-dependent^8-10,18^ or - independent^11,43^ switch of cardiomyocyte metabolism from oxidative phosphorylation to glycolysis, we next sought to challenge our findings by modulating glycolysis during heart regeneration, irrespective of HIF-1α levels. To that end, one-day old mice were randomized to sham or cryoinjury operation, the latter followed by daily subcutaneous injections of PBS (control) and either a low- (2.5mg/kg) or high-dose (100mg/kg) of metformin, for a total of 7 days. Metformin is a glucose-lowering drug with pleiotropic, incompletely understood mechanisms of action, involving reducing gluconeogenesis through activation of AMPK, inhibition of cAMP and complex I of the respiratory chain; as well as enhancing insulin sensitivity and glucose uptake through the regulation of somatotrope, corticotrope, and gonadotrope functions^44,45^. Recent work indicates that metformin exerts a direct, AMPK-dependent biphasic effect on cardiomyocytes; at lower doses it enhances oxidative phosphorylation while at higher doses it induces a glycolytic switch^46^. Consistently, treatment of regenerating neonatal mice with metformin enhanced cardiac AMPK activation and resulted in lower blood glucose and higher lactate levels in the high-dose group but not in the mice that received the lower dose [Suppl. Fig. 4D-I]. Moreover, the binary effects of metformin on glycolysis corresponded well with changes in HIF-1α activity. Specifically, HIF-1α was downregulated in the low-dose group, compared to controls [Suppl. Fig. 4H-J]. Intriguingly, consistent with previous work on the pituitary gland of non-human primates^44^, metformin activated somatotrope-related mechanisms in a dose-dependent manner, as indicated by the increase in GHRH receptor (GHRH-R) cardiac protein expression compared to the control treatments [Fig. 5A-B]. Collectively, these data demonstrate that glucose metabolism could be experimentally modulated in the regenerating neonatal mouse heart by metformin and that metformin’s binary cardiac effects are linked to the activation of both AMPK and GHRH/GHRH-R signaling pathways.

**Figure 5.**
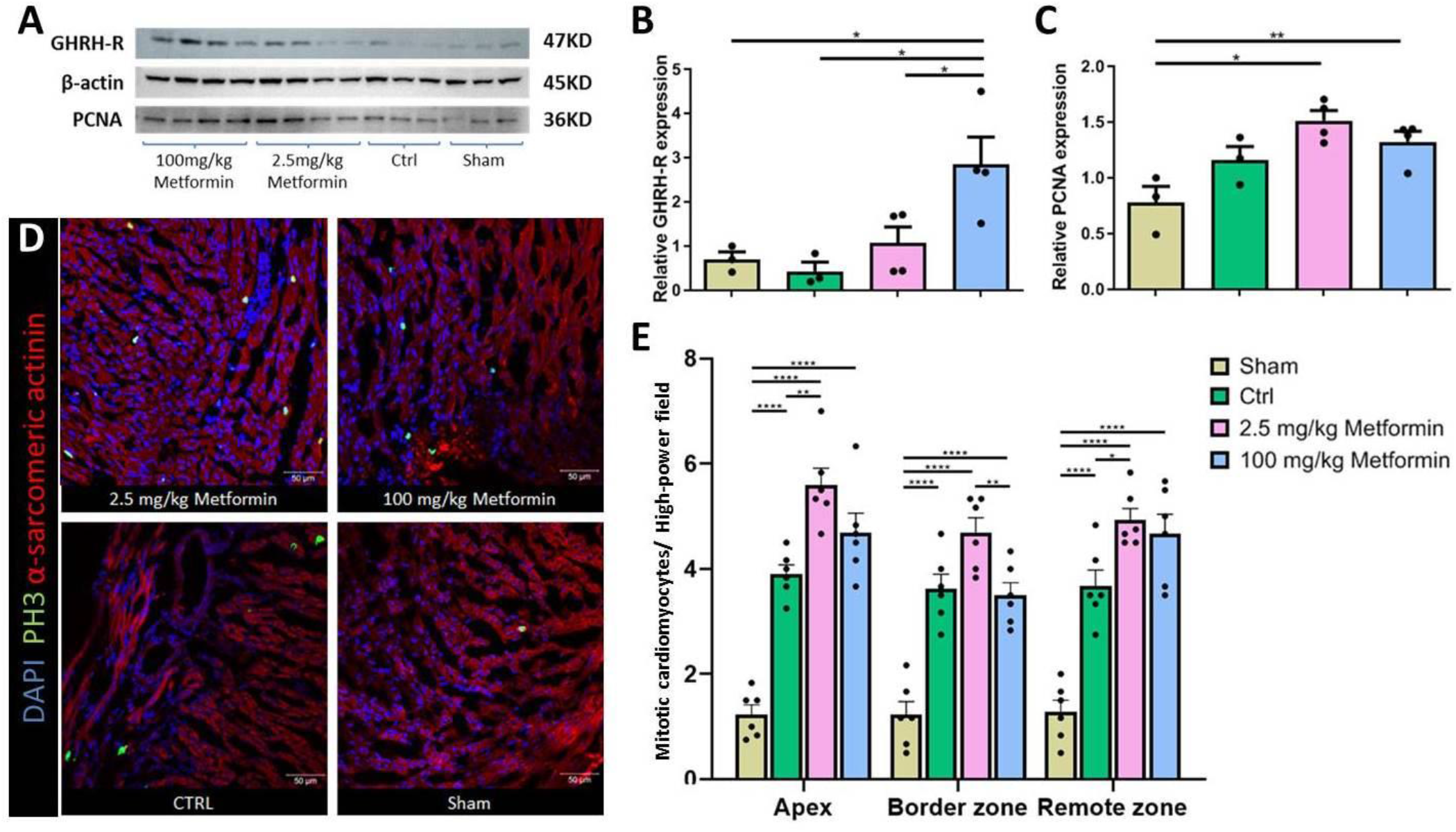
Metformin affects cardiomyocyte proliferation after cryoinjury in a dose-dependent manner. **(A)**, Western blot analysis of GHRH-R and PCNA expression in day-7 post-cryoinjury hearts. **(B)**, metformin induces GHRH-R expression in a dose-dependent manner. **(C-E)**, low-dose metformin increases cardiomyocyte proliferative capacity 7 days post-cryoinjury as indicated by estimation of PCNA (**C**) and PH3 levels (**D-E)**. However, this effect is abrogated in the high-dose metformin group, suggesting that switching cardiac regenerative metabolism to glycolysis inhibits rather than enhancing cardiomyocyte mitosis. Scale bar, 50μm. (***p≤0.001, **p≤0.01, *p≤0.05) (n=6/group).

Analysis of cell cycle activity as measured by the expression of proliferating cell nuclear antigen (PCNA) (western blotting) and PH3 (confocal immunofluorescence) showed that treatment with low-dose metformin augmented the proliferative capacity of day 7 post-injury regenerative cardiomyocytes [Fig. 5C-E]. On the other hand, these effects on day 7 post injury cardiomyocytes were abrogated upon induction of glycolysis in the high-dose metformin group.

Taken together, these results parallel our findings from the hiPSC-based cardiomyocyte differentiation studies suggesting a rate-limiting role of HIF-1α-induced aerobic glycolysis in cardiomyocyte proliferation. Both sets of experiments demonstrate that inhibition of glycolytic rewiring enhances the proliferative capacity of regenerative neonatal mouse cardiomyocytes and hiPSC-derived cardiomyocytes.

### Oxygen-independent regulation of cardiac HIF-1α through GHRH/GHRH-R

GHRH is secreted from the hypothalamus and acts on somatotropes through GHRH-R to regulate the activity of the GH/IGF-1 axis. During fetal and postnatal growth, the activity of GHRH itself is regulated through steroid sex hormones^47^. The dose dependent increase in GHRH-R expression in response to metformin [Fig. 5A-B], suggests that GHRH signaling could confer cardiomyocyte-specific control of glycolytic metabolism and, thereby, aerobic control of HIF-1α in the developing and regenerating mammalian heart. Therefore, the roles of GHRH/GHRH-R in the developing and postnatal heart were investigated further.

Gene expression analysis in the hiPSC model of human cardiomyogenesis [Fig. 6A] showed that both GHRH and GHRH-R expression increased over the time course of differentiation consistent with increased expression of the bHLH master cardiac transcription factor NKX2-5 [Fig. 6B]. Confocal and flow cytometric analyses confirmed that GHRH-R is expressed on the surface of most, if not all, NKX2-5 expressing cardiomyoblasts, both before and after their differentiation into cardiac troponin-T^+^ beating cardiomyocytes [Fig. 6C, Suppl. Fig. 5A]. Moreover, co-localization with the LIM homeodomain transcription factor ISL1, indicated that GHRH-R is expressed in both left- and right-sided cardiomyoblasts [Fig. 6C]. Similarly, the NKX2-5 cardiomyoblast-specific GHRH-R expression was confirmed through immunohistochemical analysis in human and mouse fetal heart tissue samples [Suppl. Fig. 5B-C]. Finally, the relationship between GHRH/GHRH-R signaling and NKX2-5 expression was genetically interrogate using a GHRH-R over-expressing hiPSC line (GHRHR-OE) generated using the CRISPR activation (CRISPRa) system [Suppl. Fig. 6A-C]^48^. Remarkably, GHRH-R over expressing cells (GHRHR-OE) exhibited more robust cardiomyocyte differentiation capacity than either the WT (dCas9-VP64 alone) or the HIF1α-KO cells [Suppl. Fig. 6D]. This increase in the cardiomyocyte differentiation capacity was accompanied by an upregulation in *NKX2-5* expression [Suppl. Fig. 6D]. Thus, these findings illustrate that during embryonic development, the growth and differentiation of NKX2-5 cardiomyoblasts is regulated through a lineage-specific, autocrine rather than endocrine, GHRH/GHRH-R signaling mechanism.

**Figure 6.**
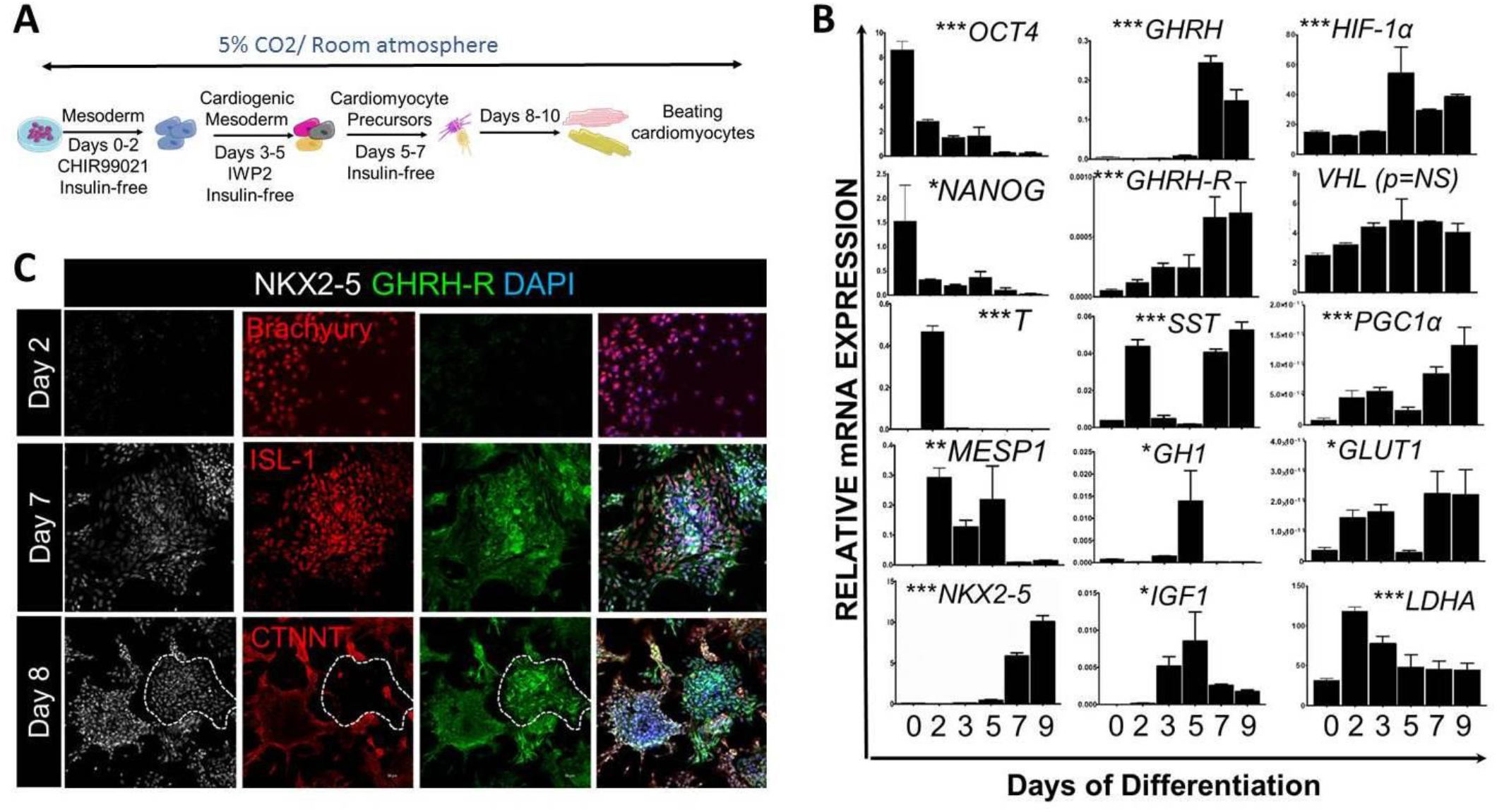
GHRH/GHRH-R signaling is an O_2_-independent regulator of cardiac HIF-1α. **(A)**, schematic illustration of the hiPSC model of human cardiomyogenesis. **(B)**, Temporal quantitative PCR analysis during cardiomyocyte differentiation indicates that *GHRH/GHRH-R* coincide with *NKX2-5* as well as *SST* expression; and are preceded by a peak in *GH1*/*IGF1* and *HIF-1α* expression. Induction of *GHRH/GHRH-R* is followed by decreased *LDHA* and increased *PGC1α* and *GLUT1* expression. **(C)**, Temporally-resolved confocal microscopy analysis of GHRH-R expression during human cardiomyogenesis. GHRH-R is not expressed in T-Brachyury^+^ early cardiogenic mesodermal progenitors but is strongly and specifically expressed on the surface of NKX2-5^+^ and ISl1^+^ hiPSC-cardiomyoblasts and cardiac troponin T^+^ beating cardiomyocytes. (***p<0.005, **p<0.005, *p<0.05) (n=3/group).

Further analyses revealed that the developmental induction of *GHRH/GHRH-R* and *NKX2-5* is preceded by a spike in *Growth Hormone 1* (*GH1*) mRNA synthesis, which in turn is preceded by a transient spike in *Insulin-like Growth Factor 1* (*IGF-1*) and *HIF-1α* transcription [Fig. 6B]. The latter findings agree with previous studies that demonstrated a reciprocal positive regulation of HIF-1α by insulin/IGF-1^49-51^. Interestingly, at later stages of differentiation, coincident with the increase in cardiomyoblast mitosis [Fig. 3B-C], the GH/IGF1/HIF-1α axis is likely antagonized by GHRH/GHRH-R, as indicated by the downregulation of *HIF-1α, GH1* and *IGF-1* mRNA synthesis and induction of the GH/IGF-1 antagonist *Somatostatin* (*SST*) from day 7 onwards [Fig. 6B]. These autocrine hormonal changes are accompanied by increased *PGC1α* and *GLUT1*, as well as a decreased *LDHA* expression [Fig. 6B], indicating a metabolic transition from non-oxidative to oxidative glucose metabolism. Finally, analysis of a publicly available microarray dataset illustrates similar spatiotemporal expression profiles during the development and postnatal growth of the mouse left ventricle [Suppl. Fig. 5D].

Importantly, HIF-1α Chip-seq analysis illustrated that both NKX2-5 and *SST* are HIF-1α target genes [Suppl. Fig. 6E], suggesting that their activation could be enhanced through the aerobic stabilization of HIF-1α. Consistent with this hypothesis, 45-min stimulation of day 7 WT hiPSC cardiomyoblasts with recombinant human (rh)GHRH led to a dose-dependent response in *SST* as well as the HIF-1α target genes *LDHA* and *GLUT1*, and these effects were blunted in HIF1α-KO cells [Suppl. Fig. 6F]. Notably, in WT cells, the expression of *SST* and *GLUT1* increased dose-dependently at lower concentrations (<1μΜ rhGHRH) and declined at higher (3μΜ rhGHRH), supporting regulation via a somatotrope-like feedback loop^52^ [Suppl. Fig. 6F].

To gain further insights into the mechanism of GHRH/GHRH-R signaling in cardiomyocyte differentiation, WT hiPSC-cardiomyocytes were stimulated for 45 min with increasing concentrations of GHRH (0μM, 0.15μM and 0.3μΜ) and the response of the cells was assessed using RNAseq analysis. ANOVA-like testing using EdgeR^53^ identified 83 DEGs between the 3 treatment groups [Fig. 7A, Suppl. Data 5]. Reactome gene set enrichment analysis found overrepresentation in terms related to HIF-α, Insulin, IGF and GH, as well as nerve growth factor (NGF, NTRKs) signaling [Fig. 7A, Suppl. Data 6].

**Figure 7.**
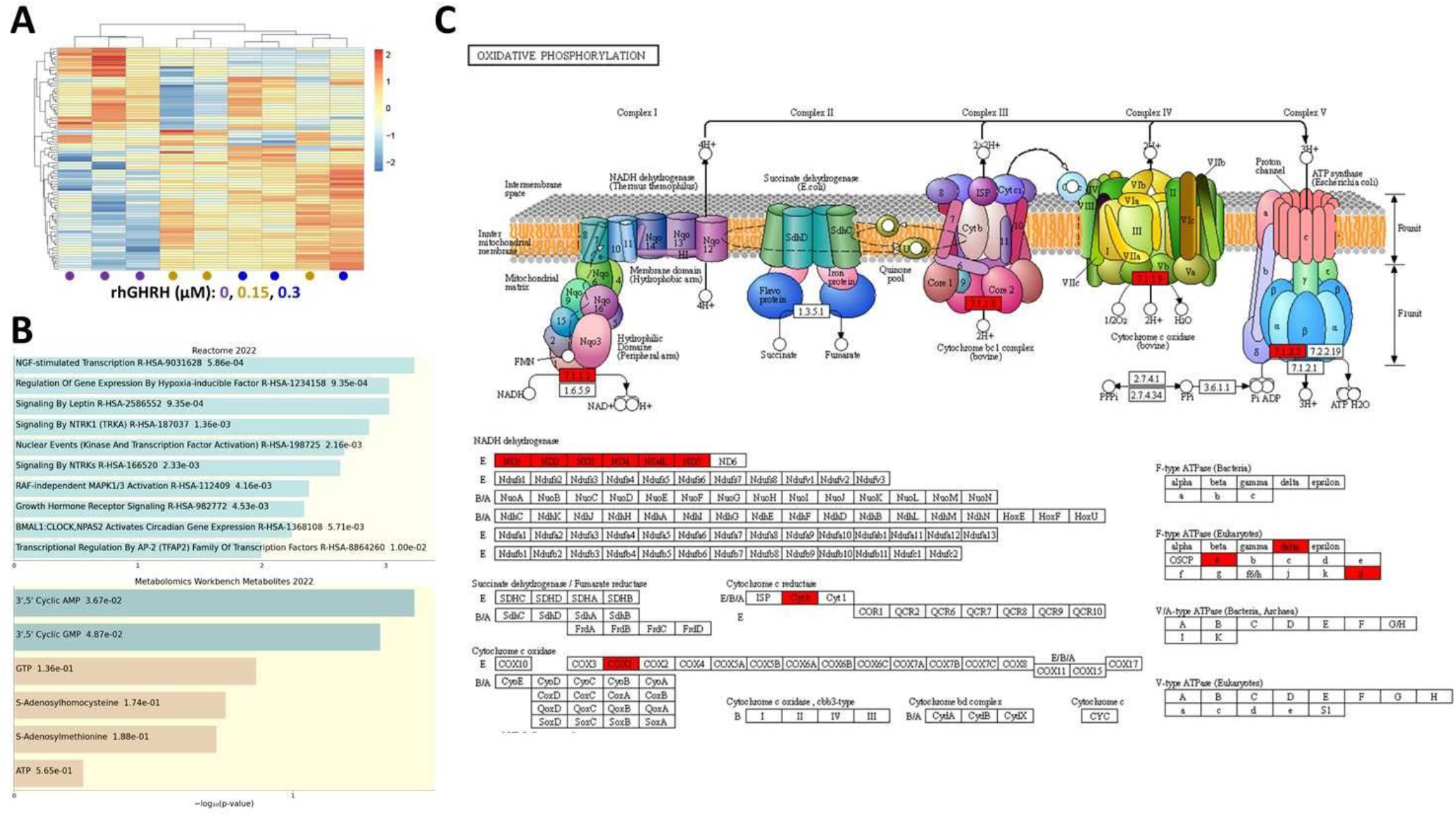
GHRH/GHRH-R signaling stimulates HIF-1α, cAMP and cGMP intracellular pathways. **(A)**, Unsupervised hierarchical clustering of the 83 differentially expressed genes in hiPSC cardiomyocytes stimulated for 45min with 0, 0.15 and 0.3μΜ recombinant human (rh)GHRH (n=3/group). **(B)**, Bar chart of top enriched terms from the Reactome_2022 (up) and Metabolomics Workbench Metabolites_2022 (down) gene set libraries. The terms are displayed based on the -log10(p-value), with the actual p-value shown next to each term. **(C)**, KEGG Oxidative Phosphorylation pathway diagram with DEGs highlighted in red.

Moreover, enrichment analysis using the Human Metabolome Gene/Protein Database (MGP) revealed significant overrepresentation in cAMP and cGMP pathways [Fig. 7B]. Finally, KEGG pathway analysis demonstrated functional enrichment in complex I, III, IV and V electron transport chain (ETC) genes [Fig. 7C, Suppl. Data 6]. Consistently, Western blot analysis revealed reduced ETC activity in response to 0.3μM rhGHRH, although seahorse analysis of hiPSC-cardiomyocyte bioenergetics did not record any changes in oxygen consumption rate, an indicator of mitochondrial respiration [Suppl. Fig. 7A-B]. However, the levels of extracellular acidification rate and total ATP were significantly increased compared to controls indicative of the induction of aerobic glycolysis in response to rhGHRH stimulation [Suppl. Fig. 7C-F]. Specifically, real time analysis of glycolysis showed that rhGHRH-treated hiPSC-cardiomyocytes exhibited higher maximum glycolysis when induced by oligomycin in the presence of glucose [Suppl. Fig. 7D], as well as higher glycolytic rate [Suppl. Fig. 7E], and glycolytic capacity [Suppl. Fig. 7F], compared to controls.

These findings demonstrate that cardiac GHRH/GHRH-R acts as a cardiomyoblast-specific signaling pathway that may have evolved as an O_2_-independent developmental mechanism for controlling HIF-1α stability and glucose metabolism and, thereby, the proliferative activity of myocardial cells in the mammalian heart.

## Discussion

This study identifies a novel mechanism for specification and proliferative growth of NKX2-5^+^ cardiac myoblasts in humans and mice revealing a coordinated O_2_-independent, hormonally regulated, HIF-1α metabolic pathway. Specifically, our work highlights a novel, NKX2-5 lineage-specific, autocrine GHRH/GHRH-R signaling pathway, which is activated in response to a GH1/IGF1/SST feedback mechanism and regulates HIF-1α in an O_2_-independent manner through intracellular activation of the second messengers cAMP and cGMP. Aerobically stabilized HIF-1α translocates to the nucleus of NKX2-5^+^ cardiomyoblasts and binds to *NKX2-5* and *SST* regulatory elements enhancing their transcription, while at the same time activating glycolytic and thyroid hormone genes, leading to overflow of lactate and Krebs cycle intermediates (Warburg metabolism) and cell cycle exit [Summary Figure]. Conversely, knockout of *HIF-1α* enhances cardiomyoblast proliferation by impeding activation of Warburg metabolism and promoting de novo fatty acid synthesis deploying citrate shuttle conversion of glucose to palmitate and stearate. Consequently, inhibition of HIF-1α or Warburg metabolism, through pharmacologic administration of the sGC/cGMP agonist YC-1 or low-dose of the cAMP inhibitor metformin, respectively, stimulates cardiomyocyte proliferation and regeneration in neonatal mice.

Our study supports previous findings in mice^12^ and quail^54^ embryos, as well as in human pluripotent stem cell-derived cardiomyocytes^55^, suggesting a repressive role for HIF-1α and hypoxia in cardiomyocyte proliferation. Furthermore, through a systems biology approach in hiPSCs and mice, we unraveled an O_2_-independent, multifaceted mechanism of action that likely underlies this effect. First, consistent with previous studies in mice and amphibians^16,56^, we showed that HIF-1α directly targets and activates the master cardiac transcription factor NKX2-5 during human cardiomyogenesis. Previous studies in mice have placed NKX2-5 at the center of the gene regulatory network that coordinates specification of ventricular myocytes into the ventricular conduction system lineage^57^. Notably, the cardiac conduction system arises from a subset of ventricular cardiomyoblasts with slowed proliferative capacity^58^; consequently, loss of NKX2-5 leads to abnormal conduction system development and ventricular cardiomyoblast hyperplasia^59^. Thus, NKX2-5 represses cardiomyoblast proliferation in favor of conduction system development; hence, its direct targeting by HIF-1α partly explains the enhanced proliferative phenotype seen in our study. Moreover, it provides a potential explanation for the increased sphericity and conduction system defects described here and elsewhere^14^, in response to myocardial HIF-1α hyperactivation by CoCl_2_ or VHL deletion, respectively.

Second, we show that increased cardiomyoblast proliferation in response to HIF-1α inactivation is fueled through a rewiring in glucose metabolism. Specifically, we found that during normal human heart development, O_2_-independent activation of HIF-1α enhances glucose oxidation to pyruvate which in turn fuels both lactate production and the Krebs cycle. More importantly, the enhanced glucose oxidation fueled leakage of Krebs cycle intermediates, including citrate, α-ketoglutarate, succinate, fumarate, and malate, which may establish a positive feedback loop that perpetuates O_2_-independent HIF-1α signaling and cell cycle inactivation^60^. Conversely, inactivation of HIF-1α rewires glucose oxidation to fuel palmitate and stearate biosynthesis, and this effect is accompanied by cell cycle re-entry [Suppl. Fig. 3C]. Several mechanisms may explain the increased cardiomyoblast proliferation in response to long-chain, saturated fatty acid biosynthesis. For instance, compared to glucose, long-chain fatty acids provide more efficient substrates both for the energy and biomass requirements of a proliferating cell^61^. In addition, several signaling pathways previously linked to cardiomyocyte proliferation, including Wnt signaling^62^, rely on palmitoylation of Wnt ligands as well as the Porcupine palmitoyltransferase. Therefore, some of the palmitate molecules may serve this purpose. Intriguingly, Wnt signaling is among the top functionally enriched terms of the non-HIF-1α target genes that are upregulated in HIF1α-ΚΟ hiPSC-cardiomyocytes [Fig. 2D].

Notably, our findings indicating redirection of glucose oxidation toward fatty acid synthesis are in line with, and may provide further mechanistic insights into, recent work demonstrating that inhibition of PDK4^63^ or succinate dehydrogenase^43^ enhances cardiomyocyte proliferation and regeneration in mice exit [Summary Figure]. Specifically, consistent with our data, rewiring of glucose metabolism from lactate synthesis toward oxidative metabolism in the Krebs cycle in response to PDK4 inhibition, enhances mouse cardiomyocyte proliferation and regeneration^63^. Intriguingly, cardiac-specific *PDK4*-knockout mice exhibited reduced exogenous fatty acid utilization as well as enhanced cardiomyocyte cell cycle progression; whereas neonatal mice fed fatty-acid deficient milk showed prolongation of the postnatal cardiomyocyte proliferative window. Although these observations may seem contradictory with our data linking enhanced cardiomyocyte proliferation to long-chain fatty acid biosynthesis, we postulate that it indicates competition between exogenous *vs*. glycolytically derived fatty acid utilization as a driver of cardiomyocyte cell cycle exit. Further studies are needed to determine whether exogenous fatty acid supplementation counteracts cardiomyocyte proliferation in HIF1α-ΚΟ hiPSC cardiomyocytes; and similarly, whether *PDK4*-knockout or fatty-acid deficient milk diet^63^ lead to increased glycolytically derived fatty acid biogenesis. Likewise, our data showing an inverse relationship between cardiomyocyte proliferation and accumulation of Krebs cycle intermediates are consistent with recent work demonstrating that inhibition of succinate accumulation through the SDH inhibitor malonate, enhances cardiomyocyte proliferation and regeneration^43^.

Importantly, our data argue against a recently proposed, paradoxical hypothesis suggesting hypoxic activation of a HIF-1α-mediated anaerobic glycolytic switch as a key driver of cardiomyocyte proliferation and regeneration in mice^10,16,42^ and zebrafish^8^. Potential explanations for this discrepancy include previously recorded inconsistencies in HIF-1α mutant mouse lines as well as evolutionary differences between the 4-chambered mammalian heart and the 2-chambered fish heart and, consequently, blood flow and oxygen supply differences between these 2 vertebrate models of heart development and regeneration^1,20^.

Finally, our work highlights the role of GHRH/GHRH-R signaling as a novel, cardiomyocyte-specific, oxygen independent regulator of HIF-1α through activation of cAMP and cGMP pathways. Interestingly, a recent cardiovascular mouse phenotyping study reported abnormal ST segments in the electrocardiograms of GHRH mutant mice^64^, suggesting inherited, non-ischemic arrhythmic disorders such as those seen in Brugada and Bundgaard syndromes^65^, or in mice with NKX2-5-specific HIF-1α activation^14^. Importantly, although our studies agree with previous work from our lab demonstrating concomitant activation of both cAMP and cGMP pathways in cardiomyocytes in response to stimulation of GHRH-R with rhGHRH or synthetic analogs of GHRH^66^, the neonatal heart regeneration experiments with the sGC/cGMP activator YC-1 and the cAMP antagonist metformin indicate a biphasic effect of cAMP and cGMP pathways on HIF-1α and cardiomyocyte proliferation [Summary Figure]. The work from our lab^47,66,67^ and others^68^ consistently demonstrate important therapeutic roles for GHRH/GHRH-R signaling stimulation in adult heart diseases. The biphasic effects of its downstream effectors cAMP and cGMP may partly explain why responses in adult cardiomyocyte proliferation and regeneration are not systematically demonstrated^47,67^. Thus, we will need to test whether a combination therapy of synthetic analogs of GHRH with e.g., low-dose metformin permits greater regenerative responses in the adult heart.

In summary, our findings demonstrate that HIF-1α signaling and induction of glycolysis suppress cardiomyocyte proliferation and regeneration in mammals and suggest that targeting GHRH/GHRH-R signaling provides a myocardial lineage-specific, O_2_-independent control of HIF-1α activity. We suggest that pharmacologic or genetic approaches that modulate this pathway could provide novel therapeutic strategies for stimulating regeneration in the adult heart.

## Author contributions

A.C.B.A.W. and A.D. performed and analyzed experiments and assisted with manuscript writing. K.P., J.N.K., S.K., T.M., D.A.R., E.I.P., G.S., P.P.S., K.K., E.T., A.G.S., K.V., and D.M.D., performed and analyzed experiments and assisted with manuscript editing. H.G.G., C.K., D.I.K., A.L., W.A.B. and A.V.S. assisted with experimental support, data interpretation and manuscript editing. J.M.H. conceived of the experimental design and assisted with experimental support and manuscript writing. K.E.H. conceived, designed, performed, analyzed and interpreted the experiments and wrote the manuscript.

## Materials and Methods

### Animal experiments

The *Mef2c-AHF-Cre;tdTomato* mouse samples have been previously described^69^. Neonatal mice cryoinjury and E11.5 embryos experiments were conducted in BALB/c mice. Animals were maintained at the Aristotle University of Thessaloniki, School of Biology. Procedures were performed according to the guidelines of the European Commission and the ARRIVE guidelines. All analyses were performed in age-matched male and female littermates from multiple litters.

### Cryoinjury surgery

The cryoinjury-induced heart regeneration mouse model was performed according to previously described protocols^39^. Briefly, 1-day-old mice were anesthetized by hypothermia and the apex of the heart was injured with a liquid-nitrogen cooled copper filament for 3 seconds. Subsequently, the thoracic wall muscles and skin were sutured, and the neonatal mice were placed on a heated surface until they regained their mobility. The same procedure was followed for the sham group, except the apex cryoinjury.

### Injections

Neonatal mice were treated with daily subcutaneous injections, from day 1 to day 6 post-injury. The following compounds were administered in a standard volume of 50μL: YC-1 (70μg in 1% DMSO; A12402, Adooq Bioscience), CoCl_2_ (120μg in saline; 232696, Sigma-Aldrich), Metformin (2.5mg/kg (low-dose) and 100mg/kg (high-dose) in saline; A10BA02, Glucoformin; Medicair), and saline. On day 7 after surgery, animals were euthanized by decapitation and the heart tissue was excised. Some pups were also injected with Hypoxyprobe (60mg/kg in saline; HP7-100, Hypoxyprobe Inc.) once, 3h before euthanasia. For embryo experiments, pregnant female mice were injected intraperitoneally with 60mg/kg Hypoxyprobe once, 3h before sacrifice and embryos excision.

### Glucose and Lactate measurement

The effects of Metformin on glucose metabolism were evaluated on 7-day-old cryoinjured mice treated with daily injections of placebo, low- or high-dose of Metformin. Mice were fasted for 24 h, then injected with Metformin or saline, and 2 hours later glucose and lactate levels were measured in peripheral blood using the Solus V2 (Eifron) and Lactate Plus (Nova Biomedical) devices, respectively.

### Human fetal heart tissue

The human fetal heart tissue samples (at 15 to 22 weeks of gestation) have been previously described and were obtained from authorized sources (Advanced Bioscience Resources Inc., Alameda, CA) following Institutional Review Board approval^69^.

### hiPSC differentiation toward cardiomyocytes

Culture and differentiation of hiPSCs (SC101A, System Biosciences) were performed as previously described, with slight modifications^70^. Briefly, hiPSCs grown in E8 medium on Matrigel-coated dishes were single-cell suspended with Tryple (GIBCO) and seeded at a density of 1×10^5^ cells/well of a 12-well plate and grown for 96h to 90% confluence in a humidified incubator with 5% CO_2_ at 37°C. On day 0, the medium was replaced with RPMI 1640 supplemented with B27 without insulin and 6 μM CHIR99021 (2520691, Biogems). On day 1, the medium was changed to RPMI 1640, supplemented with B27 without insulin for 72h, supplemented with 5μM IWP2 (6866167, Biogems) after the first 24h. On Day 4, the medium was switched to RPMI 1640 supplemented with 3μg/mL heparin until day 8, with a medium change every 2 days. From that day on, fresh RPMI 1640 with 3μg/mL heparin and 20μg/mL insulin was changed every two days.

### Immunohistochemistry and immunocytochemistry

For immunocytochemical analysis, cells were fixed in 4% paraformaldehyde, blocked for 1 hour with 10% normal donkey serum or 2% BSA, and incubated with primary antibody overnight at 4°C. Next, the cells were incubated with an Alexa Fluor-conjugated secondary antibody (1:500; Thermo Fisher Scientific) for 1h at RT, stained with fluoroshield-DAPI mounting medium and visualized on a Zeiss LSM 780 confocal microscope. Accordingly, immunofluorescence analysis of mouse heart tissues was performed in 5-to 10-µm-thick cryosections. For HIF-1α, the signal was enhanced by incubating the sections in biotinylated anti-rabbit IgG antibody (1:500; BA1000, Vector Laboratories) for 2h, followed by incubation in streptavidin-Alexa Fluor 488 dye (1:1000; S32354, Life Technologies) for 1h. For Hypoxyprobe injected mice and embryos, cryosections were incubated overnight with Hypoxyprobe Red Mab 549 (1:200; HP7-100, Hypoxyprobe Inc.). Immunofluorescence analysis of human fetal heart samples was performed in 4-to 5-µm-thick formalin fixed paraffin-embedded tissue sections. Antigen unmasking was performed by microwaving the slides for 2 × 10 min in citrate buffer solution (pH 6) (Thermo Fisher Scientific). Sections were then blocked for 1 hour at room temperature with 10% normal donkey serum (Chemicon International Inc., Temecula, CA), followed by overnight incubation at 4°C with the primary antibody. The following primary antibodies were used: HIF-1α (1:100; NB100-479, Novus Biologicals); sarcomeric α-actinin (1:400; ab9465, Abcam); PH3-Alexa Fluor488-conjugated (1:3000; ab200614, Abcam); T-Brachyury (1:50; AF2085, R&D systems); ISL-1 (a mixture of 1:10 of #40.2D6 and 1:100 of #39.4D5, Developmental Studies Hybridoma Bank); cardiac troponin-T (1:200; ab83774, Abcam); GHRH-R (1:500; ab28692, Abcam); NKX2-5 (1:50; Santa Cruz Biotechnology, SC8697).

### Western blotting

Whole cell and tissue protein was extracted using the Active Motif protein extraction Kit and quantified with the Bradford assay (Bio-Rad). Snap-frozen cardiac apical tissues were homogenized using Dounce tissue grinder (Sigma-Aldrich). Electrophoresis was performed in precast, NuPage 4 to 12% bis-tris protein gels (NP0323, Thermo Scientific; or MP42G15; Merck) before transferring into 0.22μm pore size polyvinylidene difluoride (PVDF) membranes (LC2002; Thermo Fisher Scientific) using the TransBlot Turbo transfer system (Bio-Rad) or the Mini Blot Module (B1000, Thermo Fisher Scientific). Western blots were performed after blocking with 3% Blotto (2325, Santa Cruz Biotechnology, Inc.) for 40’, using antibodies against: HIF-1α (1:2000; D1S7W, Cell Signaling), GHRH-R (1:500; ab76263, Abcam for mouse tissue), GHRH-R (1:1000; LS-C383690, LSBio, for human cells) β-actin (1:6000; 8H10D10, Cell Signaling), PCNA (1:2000; D3H8P, Cell Signaling), Gapdh (1:2000; D16H11, Cell Signaling Technology), Cas9 (1:2000; 14697, Cell signaling), AMPKα (1:2000; 5831S, Cell signaling), Thr172-phospho-AMPKα (1:2000; 2535S, Cell signaling), Total OXPHOS WB Antibody Cocktail (ab110413, Abcam), anti-Rabbit and anti-Mouse IgG, HRP-linked secondary antibodies (1:500; 7074 and 7076, Cell Signaling). The blots were visualized using the iBright imaging system (Thermo Fisher Scientific) and densitometry analysis was performed on Fiji ImageJ.

### Flow Cytometry

Flow cytometry analysis was performed on different stages of hiPSC cardiomyocyte differentiation. Cells were collected using TrypLE (12604-021, GIBCO) and fixed in 70% methanol. For cell cycle analysis, cells were incubated with Fx-Cycle PI/RNase Staining Solution (F10797, Thermo Fisher Scientific) for 30’ in room temperature, and analyzed in a BD Accuri C6 Plus cytometer. Flow cytometry data analysis was performed on FlowJo 10.8.0. In addition, the following antibodies were used in day-7 hiPSC-derived cardiomyoblasts, which were analyzed in a BD LSR-II cytometer: Troponin (BS10648-PE, Bioss), GHRH-R (C717859, LsBio), NKX2-5 (orb-103103, Biorbyt).

### Quantitative PCR

qPCR analyses were performed as described before^69,70^. Briefly, following total RNA extraction with the RNeasy mini kit (74106, Qiagen) and complementary DNA synthesis with the high-capacity cDNA reverse-transcription kit (4368814, Applied Biosystems), the samples were subjected to quantitative PCR in an iQ5 real-time PCR detection system (Bio-Rad), using the TaqMan Universal Master mix (Applied Biosystems). The following probes were used: TBP (Hs00427621), T-Brachyury (Hs00610080), Mesp1 (Hs00251489), Nkx2-5 (Hs00231763), Oct4 (Hs04260367), Nanog (Hs02387400), GHRH-R (Hs0181591), GHRH (Hs00184139), HIF-1α (Hs00153153), Gh1 (Hs00236859), SST (Hs00356144), LDHA (Hs01378790), VHL (Hs00184451), IGF1 (Hs01547656), SLC2A1 (Hs00892681), PPARGC1 (Hs00173304).

### CRISPR/Cas-mediated gene knockout and activation

The hiPSC lines stably expressing spCas9 and dCas9-VP64 were generated by transduction with lentiCas9-Blast (52962, Addgene) and lenti-dCas9-VP64-Blast (61425, Addgene) lentiviral particles, as described before^22,48^. For HIF-1α knock-out, sgRNA vectors targeting TTCACACATACAATGCACTG and GATGGTAAGCCTCATCACAG of the human HIF-1α gene (ENSG00000100644) were cloned into the pLH-spsgRNA2 vector (64114, Addgene). For GHRH-R activation, sgRNA vectors targeting TGTCAGGGGACAGCAGGGGA and AGCAGAGGGTGCGGTGGAAA of the human GHRHR gene (ENSG00000106128) were cloned into the lenti-sgRNA (MS2) puro backbone vector (73795, Addgene). Lentiviral particles were generated by transient co-transfection with the pMD2.G (12259, Addgene) and psPAX2 (12260, Addgene) plasmids into H293TN cells, using the jetPRIME kit (Polyplus), according to manufacturer’s instructions. At 48 hours following lentiviral transduction, hiPSCs were selected with hygromycin B (50μg/ml; 843555001, Roche Diagnostics), puromycin (1μg/ml; A11138-03, GIBCO) or blasticidin (10μg/ml; A11139-03, GIBCO) for a total period of 5-7 days.

### RNA sequencing

RNA was isolated with RNeasy plus mini kit (Qiagen) according to the manufacturer’s protocol. Libraries were prepared and sequenced by the Oncogenomics Shared Resource at the University of Miami, on the Illumina NextSeq 500 (single-end 75 base pair sequencing). Next generation sequencing quality was assessed using FastQC (v0.11.3). Reads were trimmed using trim galore, aligned to the human genome build hg38/GRCH38 using STAR^71^, and counts were generated using RSEM^72^. Differential expression analysis was performed with EdgeR^53^. A p-adjusted value <0.05 was used as cut-off criteria.

### HIF-1α ChIP-seq

HIF1α-bound chromatin immunoprecipitation, sequencing and bioinformatics (ChIP-seq) were performed on hiPSC-CMs, according to previously described protocols^73^. Briefly, chromatin was sonicated to an average fragment size of 200 base pairs (bp) with a Covaris M220 sonicator before ChIP. A total of 5μL of HIF-1alpha antibody (39665, Active motif) was used for each ChIP (n=2). Libraries were prepared using the NEBNext Ultra kit and sequenced on the Illumina NextSeq 500 (1 × 75 bp). Peak-calling on the ChIP-seq datasets was performed with MACS2 and the resulting peaks with a q-value ≤ 1×10^−100^ were annotated with genomic context information using ChIPpeakAnno^74^. Motif discovery analysis of the DNA sequences ±50bp from the peak summits was performed with MEME^74^.

### Seahorse real-time metabolic characterization

The mitochondrial oxygen consumption rate (OCR, O2 mpH/min) and extracellular acidification rate (ECAR, mpH/min) of hiPSCs were analyzed by a Seahorse XFe96 extracellular flux analyzer (Agilent technologies). For OCR studies, hiPSCs were kept in OCR medium (RPMI base, 25mM glucose, 1mM pyruvate, 2mM L-glutamine [pH 7.35]), and were analyzed using the Mito Stress Kit (103015-100, Agilent technologies). Cells were plated in 96-well flat-bottom Matrigel-treated plates and incubated in a non-CO_2_ incubator for 1h at 37°C. The ATP-linked respiration was quantified by subtracting the proton leak to the basal OCR. The total cellular ATP production rate is the sum of glycolytic and mitochondrial ATP (ATP Rate assay Kit; 103592-100, Agilent technologies). For ECAR analysis, hiPSCs were kept in ECAR medium (DMEM base [no bicarbonate] with 2mM L-glutamine, 143mM NaCl, and 0.5% phenol red [pH 7.35]). The complete ECAR analysis (Glycolysis Stress Kit; 103017-100, Agilent technologies) consisted of four stages: basal (without drugs), glycolysis induction (10mM glucose), maximal glycolysis induction (2mM oligomycin), and glycolysis inhibition (100mM 2-DG).

### Metabolomics GC-MS analysis

Exometabolites were extracted from 24-hour incubated day-0 hiPSCs or day-10 differentiated cardiomyocytes by saving the spent medium in -80°C. For the endometabolome extraction, the cells were washed with serum, followed by incubation with 100% MeOH in -20°C for 20’. Next, cells were scraped and centrifuged in 5000rpm for 15’, and the metabolites pellet was stored in -80°C until analyzed.

For GC-MS analysis, 10 μL of myristic acid-d27 (100 μg/mL; 60658-41-5, Sigma-Aldrich) and 10 μL 4-phenylbutyric acid (100 μg/mL; 1821-12-1, Sigma-Aldrich) were added to 140 µL of intracellular and extracellular extracts followed by evaporation to dryness under vacuum (Speedvac Eppendorf). The dried residues were reconstituted in 50 µL of 2% MeOX in anhydrous pyridine (110-86-1, Sigma-Aldrich) followed by derivatization at 70°C for 2 hours. After completion of the first reaction, the samples were cooled to room temperature. Then, 100 μL MSTFA 1% TMCS (24589-78-4, Sigma-Aldrich) were added and the second derivatization was performed for 1 h at 70°C. Ten (10) μL of injection standard (N-pentadecane, 100 μg/mL; 629-62-9, Sigma-Aldrich) were added and the samples were split into 2 vials for targeted and untargeted analysis.

GC-MS analysis was performed on an EVOQ 456 GC-TQ-MS system (Bruker, Billerica, MA, USA) equipped with a CTC automatic sampler and PTV injector, controlled by Compass Hystar software. A 30 m HP-5 MS UI (Agilent J&W) column (0.25 mm, ID of 0.25 μm) was used. Analytical conditions are provided in more detail previously^75^. Data were processed with AMDIS software used to achieve chromatographic peak deconvolution and identification. NIST (mainlib) and FIEHN libraries were used for peak identification. Peak areas of the compounds extracted by AMDIS were calculated using the Gavin3 script in MATLAB.

### Statistical analysis

All statistical tests were performed in GraphPad Prism (version 8, La Jolla, CA) using Student’s t test, Mann-Whitney test, or one-way analysis of variance (ANOVA) followed by Tukey’s post hoc tests. All data met the assumptions of the tests (Bartlett’s test for normality). A *p* value of <0.05 was considered statistically significant. All values are reported as means ± SEM.

## Supplementary Figure Legends

**Figure S1.**
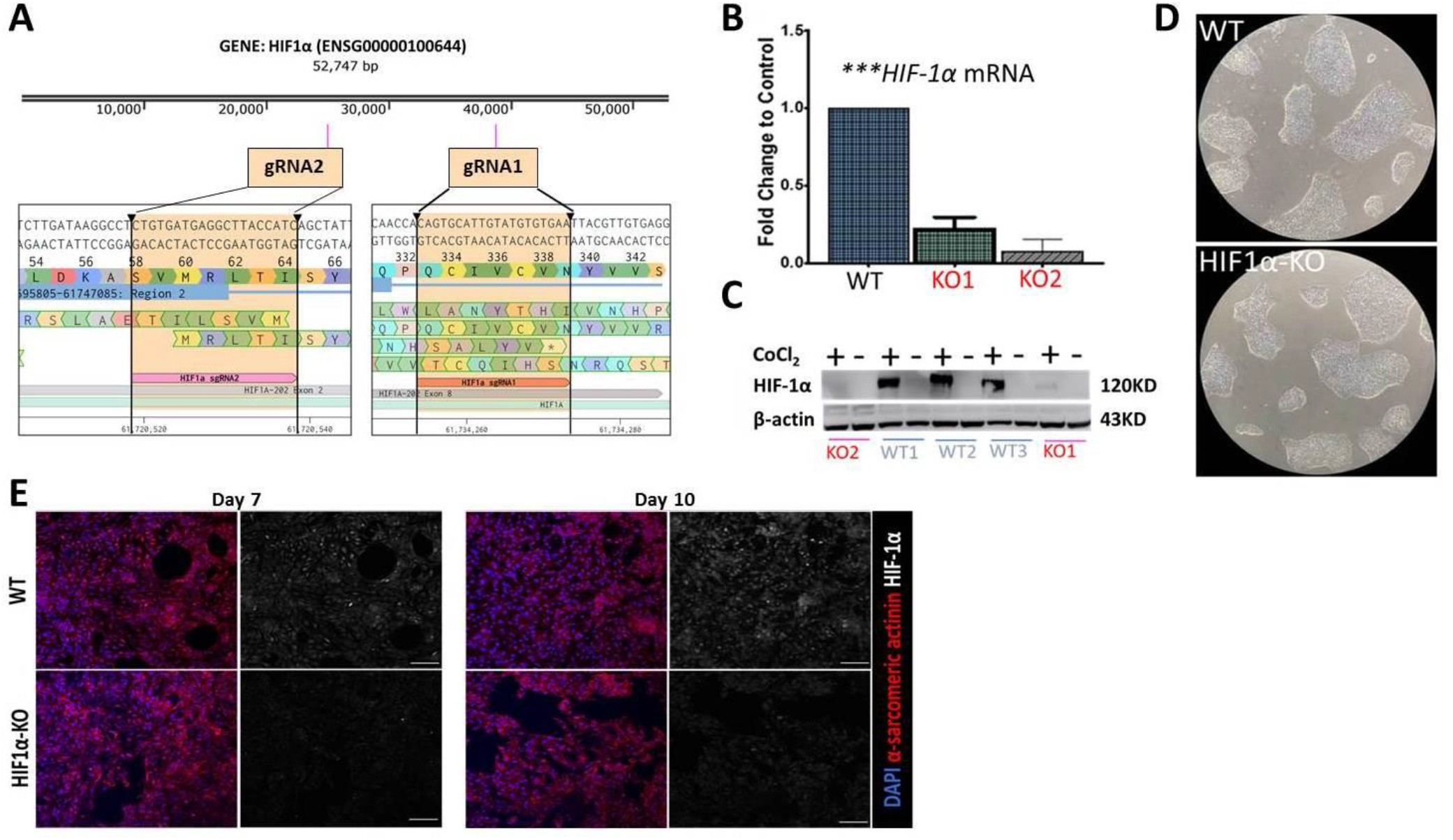
Knockout of HIF-1α does not impact the renewal and cardiomyocyte differentiation of hiPSCs. **(A)**, schematic illustration of *HIF-1α* gene editing in two distinct regions through CRISPR-Cas9. **(B-C)**, qPCR and western blot analysis demonstrate successful generation of two HIF1α-ΚΟ hiPSCs lines. (**D)**, WT and HIF1α-KO hiPSCs are phenotypically indistinguishable. (**E)**, confocal microscopy demonstrating robust nuclear HIF-1α immunoreactivity in WT (upper panels) but not HIF1α-KO (lower panels) hiPSCs, on days 7 (left) and 10 (right) of cardiomyocyte differentiation. Scale bars, 50μm.

**Figure S2.**
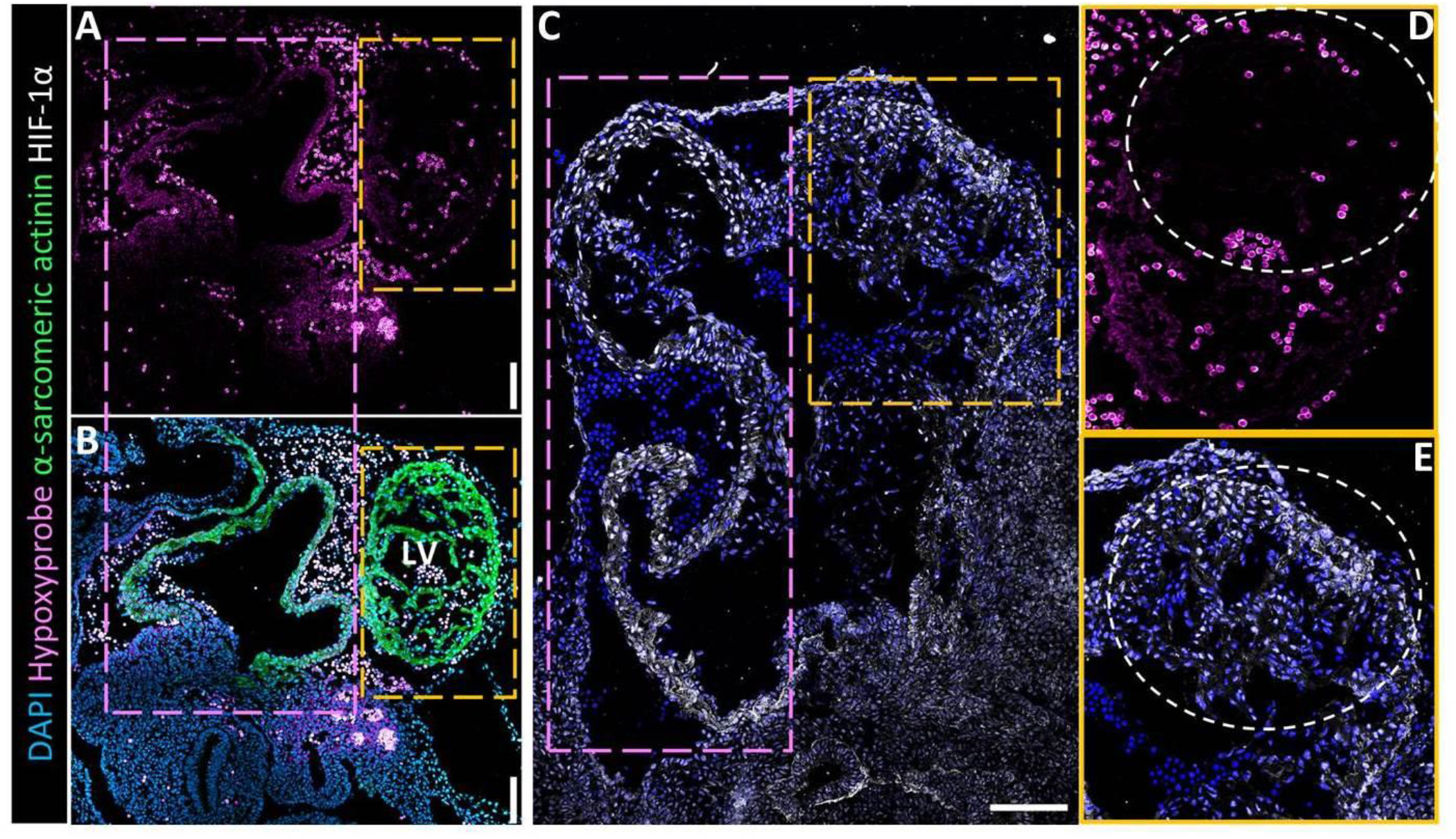
HIF-1α is stabilized in an oxygen-independent manner in the developing mouse heart. **(A-B)**, Confocal microscopy on hypoxyprobe-treated, E11.5 mouse embryos demonstrating both the presence (magenta boxed areas) and lack (yellow boxed areas) of hypoxia in the developing mouse heart. **(C)**, An adjacent cryosection to **A-B**, stained for HIF-1α, demonstrates strong nuclear HIF-1α immunoreactivity in both the hypoxic (magenta box) and non-hypoxic (yellow box) myocardium, indicating oxygen-independent stabilization of HIF-1α. **(D-E)**, Magnification of the yellow-boxed area in **C**. The white-circled area demarcates hypoxyprobe-negative left ventricular myocardium exhibiting strong nuclear HIF-1α immunoreactivity. LV: left ventricle. Scale bars, 100μm. (n=3).

**Figure S3.**
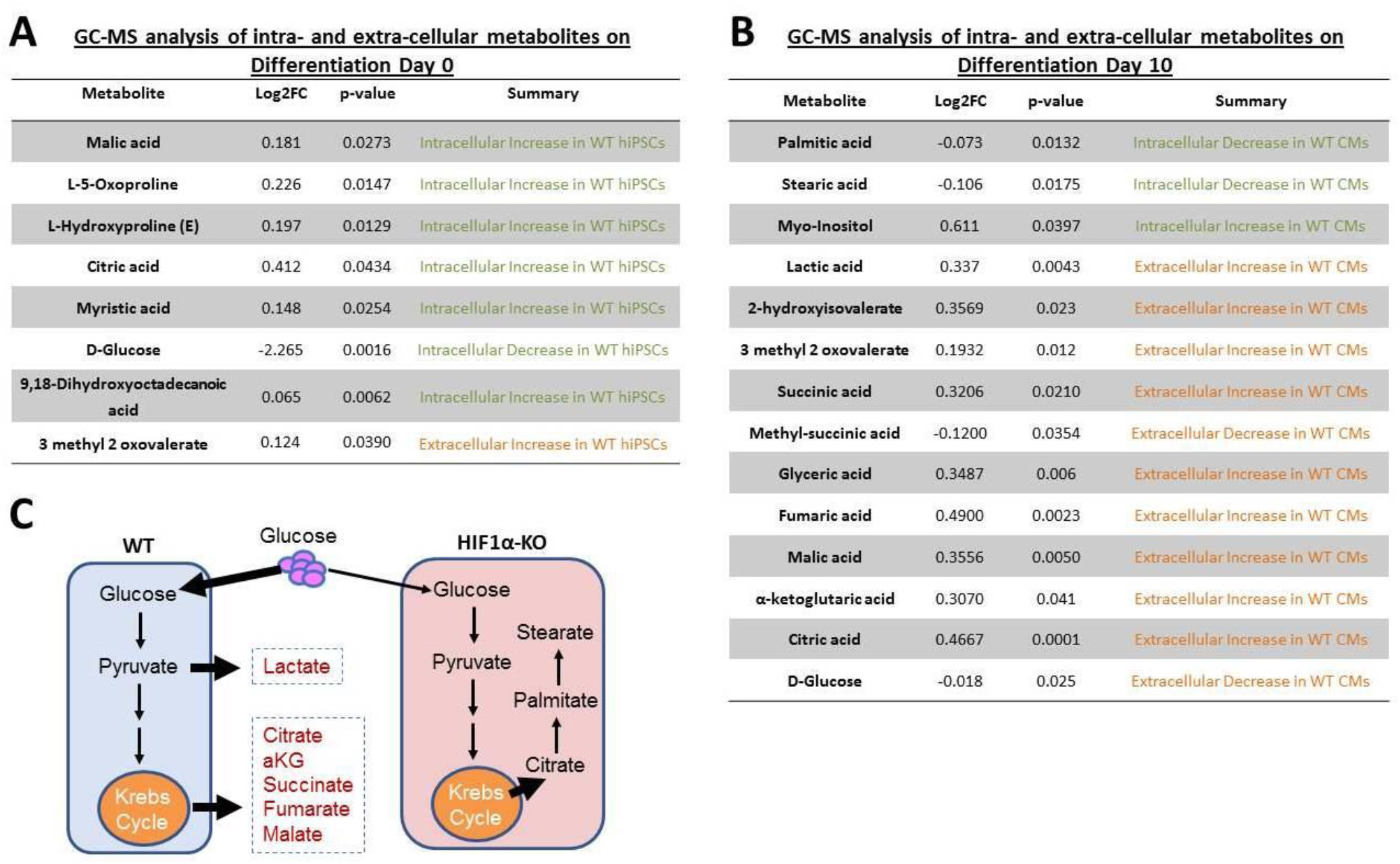
HIF-1α induces aerobic glycolysis during human cardiomyogenesis. **(A)**, GC/MS analysis of intracellular (green) and extracellular (orange) metabolites on day 0 of cardiac differentiation. Decreased intracellular levels of D-Glucose and increased levels of Krebs cycle intermediates (i.e. citrate and malate) in WT vs HIF1α-ΚΟ hiPSCs, suggest impaired aerobic glucose metabolism (Warburg effect) in HIF1α-ΚΟ hiPSCs. **(B)**, On day 10, WT hiPSC cardiomyocytes exhibit increased extracellular lactic acid and decreased glucose concentration as well as accumulation of Krebs cycle intermediates (i.e. fumarate, citrate, 2-ketoglutarate, malate, and succinate); whereas HIF1α-ΚΟ hiPSC cardiomyocytes exhibit increased intracellular levels of palmitate and stearate, suggesting that HIF-1α Knockout redirects glycolytically-derived citrate toward saturated fatty acid biosynthesis. **(C)**, Graphic summary of metabolomic studies on day 10 hiPSC-cardiomyocytes. (n=3 /group).

**Figure S4.**
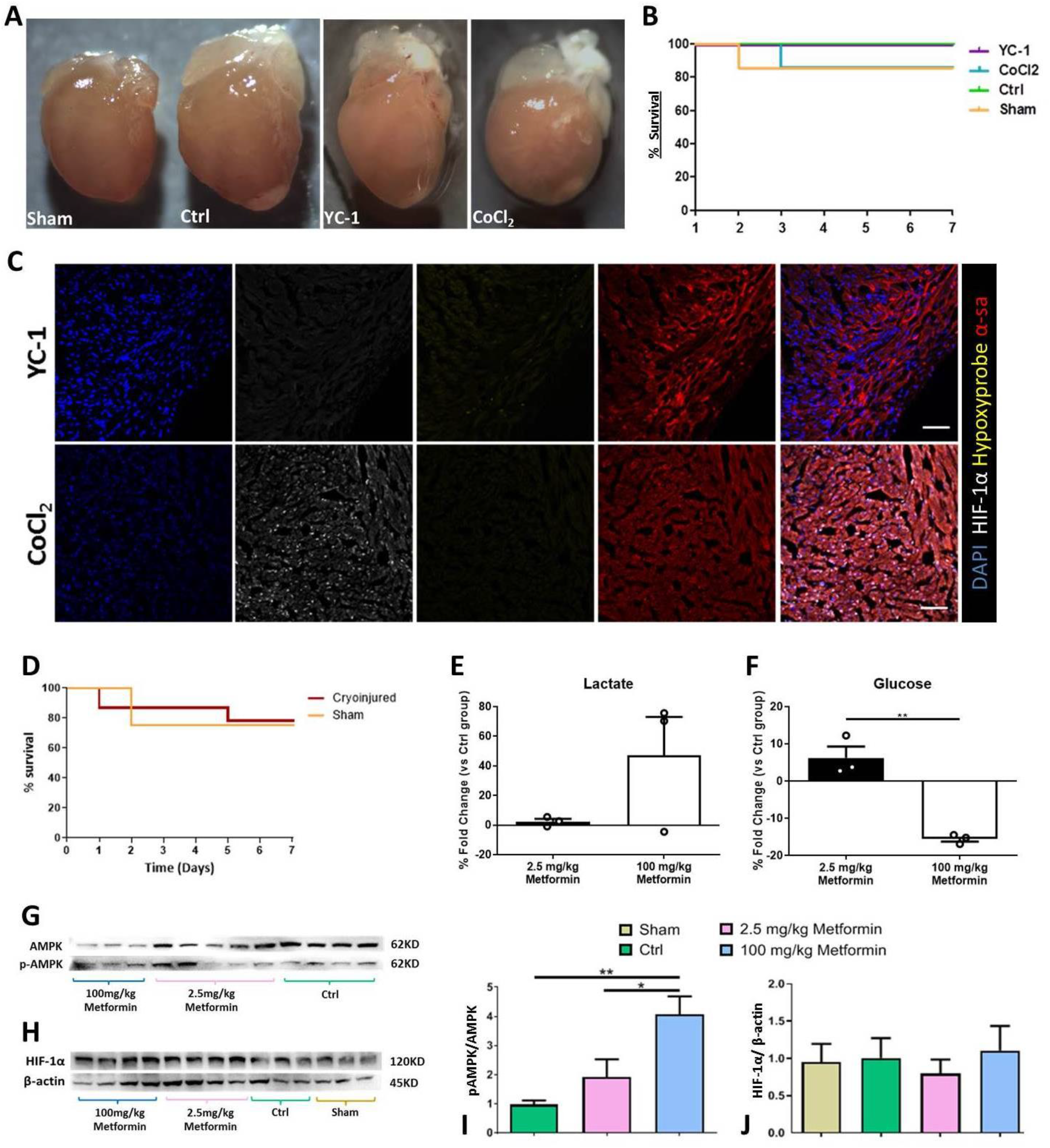
Pharmacologic modulation of HIF-1α or glycolysis in neonatal mouse hearts after cryoinjury. **(A)**, Representative macroscopic images of Sham, placebo (ctrl), YC-1, and CoCl_2_ –treated mouse hearts 7 days post operation. Tissue decoloration indicates the presence of scar tissue at the apex of the cryoinjured, but not sham-operated, hearts. Note the increased sphericity in CoCl_2_–treated heart. **(B)**, Kaplan Meier curve showing the survival rate of the experimental animals based on the compound administered in each study group (p>0.05). **(C)**, confocal microscopy on 7-day-old mice demonstrates that YC-1 (upper panels) successfully inhibited HIF-1α expression, while CoCl_2_ (lower panels) induced strong nuclear HIF-1α immunoreactivity in most cardiomyocytes, in the absence of hypoxia, as indicated by the lack of Hypoxyprobe signal. α-sa, α-sarcomeric actinin. Scale bars, 50μm. (n=6/group). **(D)**, Kaplan Meier curve showing the effectiveness of the surgical cryoinjury process. **(E-F)**, Lactate and glucose levels in peripheral blood of 24-hr fasted animals, 7 days after cryoinjury and daily low- or high-dose metformin injections. Values are depicted as % change relative to placebo-treated cryoinjured mice (n=7-8/group) (\*\**p*≤0.005). **G**, Western blot analysis of phosphorylated (pAMPK) and total AMPK. **H**, Western blot analysis of HIF-1α and β-actin expression. **I-J**, Densitometric quantification of D and E western blots, respectively (\**p*=0.05 and \*\**p*=0.01).

**Figure S5.**
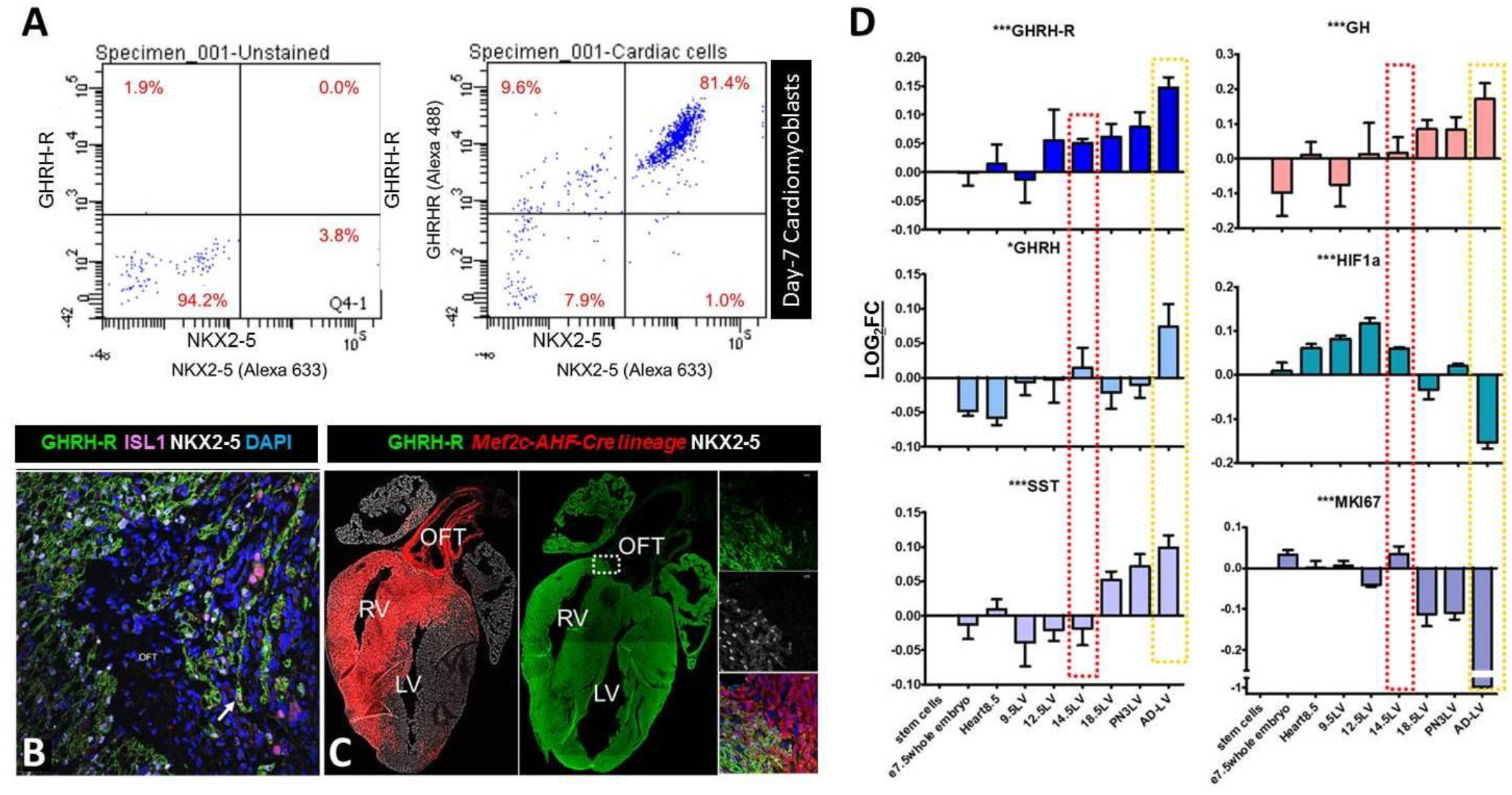
Expression of GHRH-R in the human and mouse heart. **(A)**, Flow cytometry analysis of GHRH-R and NKX2-5 co-expression on day-7 hiPSC-derived cardiomyoblasts. **(B-C)**, Confocal immunohistochemistry demonstrates that GHRH-R is specifically expressed on the surface of NKX2-5 and ISL1 (B, arrow) cardiomyoblasts, both in the developing human (**B**), and mouse (**C**) hearts (n=2). Analysis with a Mef2C-AHF-Cre reporter mouse line further demonstrates that GHRH-R is expressed in both right- and left-sided atrioventricular cardiomyocytes (**C**). **(D)**, Gene expression analysis of left ventricular samples from GSE51483, a publicly available dataset of mouse cardiac development and postnatal growth. Consistent with the hiPSC data, GHRH/GHRH-R expression is first detected in the heart at embryonic day (e) 8.5. A spike in the proliferation marker Mki67 is shown at e14.5, which coincides with increased GH and GHRH/GHRH-R signaling and decreased SST and HIF-1α. In contrast, inhibition of proliferation during the perinatal and adult stages coincides with maximum GH and GHRH/GHRH-R antagonism by SST. (***p<0.005, **p<0.005, *p<0.05) (n=3/group).

**Figure S6.**
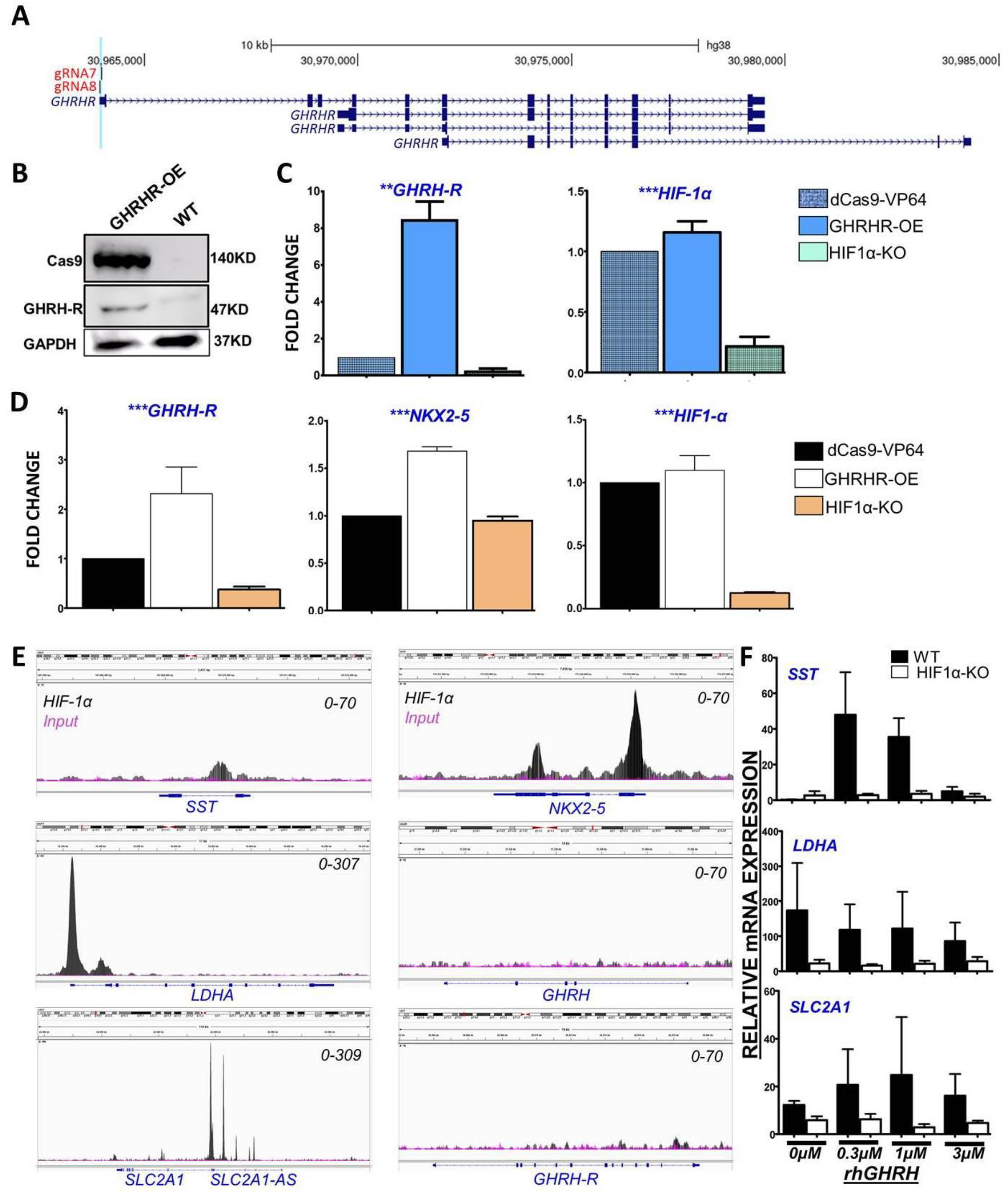
GHRH/GHRH-R signaling dose-dependently affects the expression of HIF-1α target genes in hiPSC-derived cardiomyoblasts. **(A)**, UCSC genome browser snapshot depicting the position of gRNAs #7 and #8 which were simultaneously expressed along with dCas9-VP64 to activate GHRH-R. **(B)**, Western blot confirming stable overexpression of GHRH-R in hiPSCs (GHRHR-OE). **(C)**, quantitative (q)PCR analysis of *GHRH-R* and HIF-1α in dCas9-VP64, GHRHR-OE and HIF1α-ΚΟ hiPSCs. **(D)**, qPCR analysis demonstrates that upregulation of GHRH-R enhances NKX2-5 expression and, consequently, cardiomyogenic differentiation, compared to HIF1α-KO and dCas9-VP64 hiPSC lines. (***p<0.005, **p<0.005, *p<0.05) (n=3/group). **(E)**, IGV genome viewer snapshots depicting HIF-1α ChIP-seq tracks (black, n=2) relative to input (magenta, n=1). *SST, LDHA, GLUT1* and *NKX2-5* are *HIF-1α* target genes, whereas *GHRH* and *GHRH-R* are not. **(F)**, qPCR analysis of HIF-1α targets in day-7 WT and HIF1α-KO cardiomyoblasts, in response to 45-min stimulation with increasing recombinant human (rh)GHRH concentrations (n=2/group).

**Figure S7.**
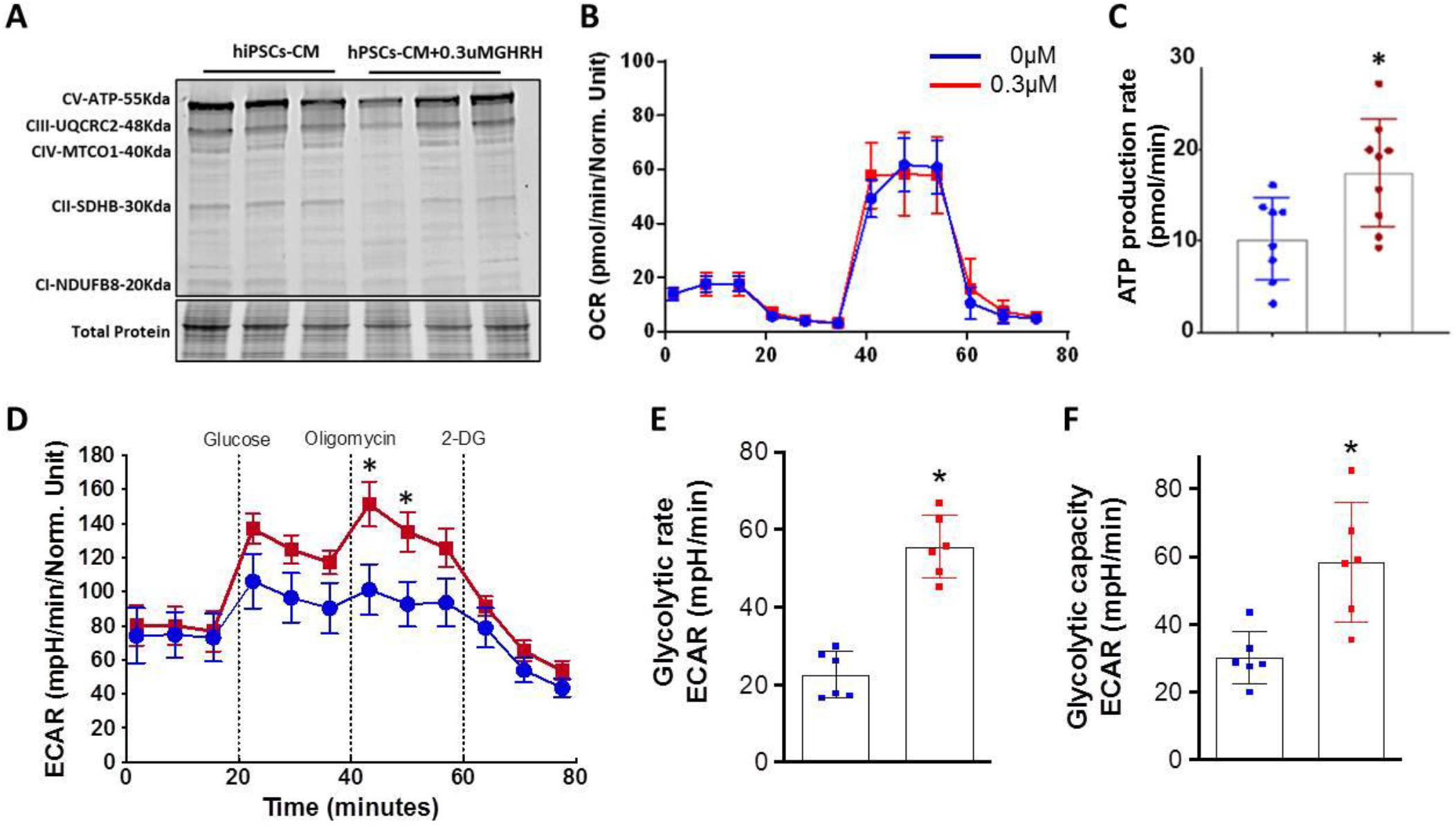
GHRH/GHRH-R activation induces metabolic changes in hiPSC-derived cardiomyocytes. **(A)**, Western blot analysis demonstrates reduced expression of mitochondrial oxidative phosphorylation complexes in response to recombinant human (rh)GHRH stimulation. **(B)**, Seahorse Live-cell Metabolic Assay measurements of oxygen consumption rate (OCR). (**C**), Total ATP production; (**D**), Extracellular acidification rate (ECAR), representing glycolysis. (**E**), Glycolytic rate (glycolysis induction subtracted for basal ECAR) and glycolytic capacity (maximal glycolysis subtracted for basal ECAR) are derived from the ECAR curve. The data were pooled from three independent experiments in duplicates or triplicates (n=6-9). *p≤0.05.

